# Partner determination from protein sequences using class information

**DOI:** 10.64898/2026.05.06.723283

**Authors:** Lisa Gennai, Francesco Caredda, Mathieu E. Rebeaud, Andrea Pagnani, Paolo De Los Rios

## Abstract

Mapping protein interaction networks and understanding how duplicated proteins acquire distinct functions require determining which paralogs from interacting protein families bind to each other.

Paralogs often fall into functional or architectural classes whose interactions can be conserved across organisms. Here, we introduce CLAPP (CLAss Pooling for Paralog matching), which pools interaction scores across organisms by class to infer one class-to-class map that constrains matching within each organism. It accepts any pairwise score assessing the likelihood of interaction of two proteins.

On nine bacterial family pairs, with test sets restricted so that the class map determines every pairing, CLAPP recovered the correct map, and hence every reference pair. Recovering the map required little information: in the two systems tested, one training epoch of a coevolution-based model sufficed, whereas an untrained model performed no better than chance; and in eight of nine systems, a conservation-only score lacking inter-protein couplings also gave perfect matching.

On unrestricted test sets, where the class map no longer uniquely determines every pairing, accuracy depended on the quality of the class labels: on 320 chaperone pairs, CLAPP performed below chance with unsupervised labels but reached 94% with phylogeny-informed ones; remaining errors involved paralogs which can interact promiscuously.

Paralog matching therefore need not rely on strong coevolutionary signals within individual pairs: when interactions between classes are conserved and labels are reliable, CLAPP can use even weak or noisy scores to accurately assign interaction partners.

## 1 Introduction

Protein-protein interactions are central to nearly all cellular processes, yet their experimental determination, while crucial, can be extremely time-consuming and resource-intensive. Computational methods stand out as a promising solution to efficiently map these interactions, guiding more targeted experimental campaigns.

As the proteome complexity of organisms increases, gene duplication and subsequent divergence leads to the emergence of paralogs, namely different proteins (or protein domains) that are present in the same organism and belong to the same protein family, typically playing different functions [1–6]. Experimental evidence indicates that paralogs in interacting protein families often evolve preferred or obligate partners [7–10]. As a result, correctly identifying which paralog pairs interact is key to understanding how interaction specificity drives functional diversification. Failure to resolve these relationships obscures the roles of individual proteins, because grouping paralogs into a single interaction class masks differences in regulation, localization, and biochemical activity that emerge after duplication. Accurate paralog matching is therefore essential for reconstructing interaction networks, assigning functional meaning to individual protein pairs, and understanding how new cellular functions evolve.

Furthermore, reliably identifying interacting paralog pairs can provide the high-quality, correctly paired datasets needed to train accurate computational models of protein-protein interactions, enabling the prediction and design of interacting protein pairs. Thus, resolving paralog matching becomes a prerequisite for fully leveraging large-scale sequence data, both to interpret existing biological systems and to support data-driven approaches to protein interaction design.

This problem is particularly relevant for molecular chaperones, where interaction specificity is closely tied to biological function. In the Hsp70 system for example, J-domain proteins (JDPs) act as co-chaperones that recruit Hsp70s to specific substrates and cellular locations, with this targeting often mediated by selective JDP–Hsp70 pairing. Both JDPs and Hsp70s have multiple paralogs within each organism, and distinct pairings have different functional roles [11]. Accurately reconstructing these interactions is therefore essential to understand how chaperone systems distribute functional tasks within the cell. Bacterial two-component signaling systems provide another example in which paralog matching is biologically informative. Sensor histidine kinases (HKs) autophosphorylate in response to specific signals and transfer the phosphoryl group to cognate response regulators (RRs), which control downstream responses. Bacterial genomes often encode many paralogous HKs and RRs, so identifying the cognate pairs can help us understand how signaling pathways retain their specificity while diversifying and avoiding detrimental crosstalk [8, 12].

Building on this need for accurate paralog matching, several strategies have been proposed. One such approach exploits genomic proximity, as interacting proteins are often encoded near each other in the genome [13–15]. However, this signal captures only part of the interaction network and is largely restricted to prokaryotic systems.

Sequence-based approaches offer a particularly attractive alternative, since interacting proteins tend to coevolve, leaving detectable signatures in their sequences. Over evolutionary timescales, shared selective constraints couple the evolution of binding interfaces and functional compatibility, so that mutations in one partner are often compensated by changes in the other. The resulting coevolutionary signal can therefore be exploited to infer interacting pairs, in the same way that coevolution has been used to reconstruct residue–residue contacts within protein families using direct coupling analysis (DCA) models [16–18]. In addition, interacting partners often display correlated evolutionary histories, as speciation and gene duplication events affecting one protein are reflected in the evolutionary tree of its interaction partner [19–21]. Comparing phylogenetic profiles or tree similarities can therefore help to identify paralog pairs that have evolved together and are more likely to interact.

Past work has formalized these ideas into a range of computational approaches for paralog matching, including phylogeny-based methods [22–32], coevolution-based frameworks [16,33–36], and later approaches combining both [37]. In parallel, protein language models have provided an alternative framework in which evolutionary, structural, and functional constraints are implicitly learned from large sequence datasets and can be used to pair interacting proteins without explicit coevolutionary modeling [38–41]. Beyond the choice of interaction score, a key practical question is how to solve the underlying one-to-one assignment between the paralogs in the interacting families. Traditional choices have been greedy or linear programming algorithms, while more recent work addresses this with a differentiable optimization framework [42].

A complementary source of information arises from the observation that many protein families can be divided into classes defined by functional, structural, or architectural features. In this work, we hypothesize that interaction specificity is often conserved at this class level across organisms, as shown in previous works for J-domain proteins and Hsp70 interactions [11, 43]. When members of one class only interact with members of another, this conserved organization provides additional constraints that can be exploited to match paralog partners. As a matter of fact, related class- or subfamily-level organization has been exploited in other paralog-matching settings [25, 28, 33, 44].

Here, we introduce CLAPP (CLAss Pooling for Paralog matching), a method that leverages this principle by pooling interaction scores across subclasses and organisms to infer paralog interactions. Rather than treating each organism independently, CLAPP aggregates evidence at the class level, reducing noise in individual interaction scores and replacing organism-specific matching problems with a shared assignment across related sequences. The framework is in principle compatible with any interaction score assessing the likelihood of interaction of two protein sequences and reduces computational cost while significantly improving accuracy. We demonstrate the performance of CLAPP on curated datasets of multiple families, including bacterial signaling systems and chaperones and co-chaperones, where it successfully recovers known interaction partners.

## 2 CLAPP Workflow

The overall CLAPP workflow consists of three main steps, illustrated in Fig. 1. First, for each organism, we assign an interaction score to every possible pair of sequences from the two protein families under study. Second, sequences within each family are assigned to classes using prior biological annotations, unsupervised classification based on sequence information alone, or a combination of both. Third, pairwise scores are pooled across organisms at the class level to infer a class-to-class interaction map, which is then used to constrain paralog matching.

**Fig. 1:**
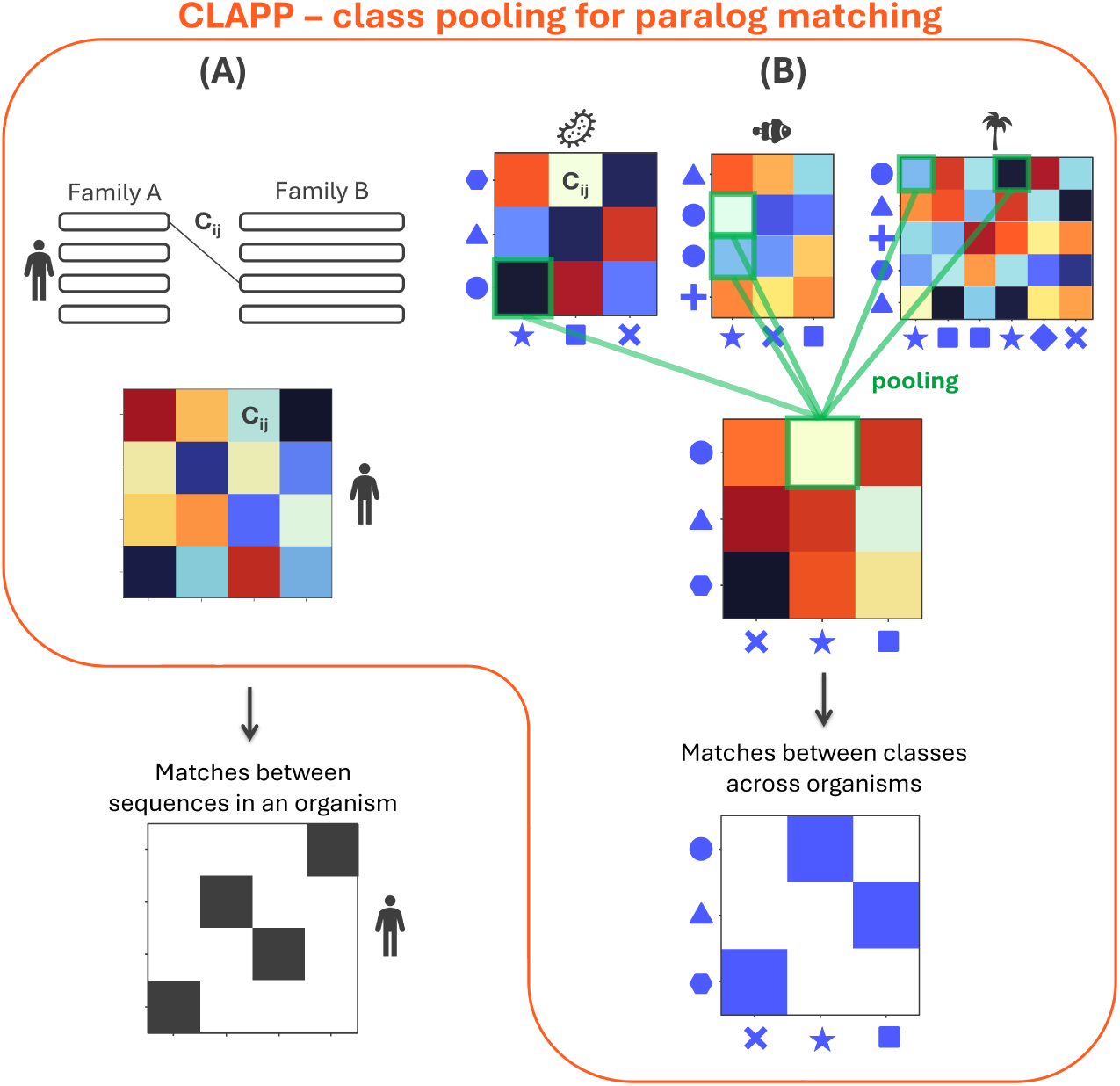
Schematic representation of CLAPP. Panel A: An interaction score is calculated for all pairs within each organism. Linear programming (LP) algorithms can also be used to find the one-to-one matches within each species that maximize the sum of the interaction scores. Panel B: pooling procedure. Interaction scores from all organisms are pooled according to the class labels of their respective sequences. Interactions between classes are then determined using LP algorithms.

More in detail, CLAPP starts from a pair of multiple sequence alignments (MSAs) of two interacting protein families. These MSAs are used to train a scoring model that assigns, within each organism, a score to every possible pair of paralogs. Here, we start with sequence-based scores that exploit the coevolutionary signal associated with protein-protein interactions. The underlying idea is that correct paralog pairs evolve under shared selective constraints and should therefore retain interaction-specific sequence signatures that distinguish them, at least statistically, from incorrect pairings.

More specifically, we score candidate pairs using a Direct Coupling Analysis (DCA) model, here AttentionDCA [45, 46], although other scoring functions could in principle be used (see section 5.1). DCA models are trained on a single MSA, which may contain sequences from an individual protein family or, as in the present case, concatenated sequences from two interacting families. The resulting score reflects the compatibility of a candidate concatenated pair with the coevolutionary patterns learned by the model.

The construction of this training MSA constitutes a key difference between CLAPP and previous paralog-matching approaches. Traditionally, such methods seek, often through iterative refinement, an optimal concatenation of sequences corresponding to the inferred interacting pairs. CLAPP instead trains on all possible within-species sequence concatenations. Correct pairs are therefore present in the training set by construction, together with many incorrect pairs, and the method relies on a crucial hypothesis: the coherent coevolutionary signal contributed by the correct pairs is sufficient to bias the learned model toward biologically meaningful interactions.

For each organism, the resulting scores define an interaction matrix containing all possible pairings between the two protein families. Standard optimization procedures, such as linear programming [47, 48] or greedy algorithms, can then be used to identify the one-to-one matching that maximizes the total interaction score.

The central additional ingredient of CLAPP is the structure of the two interacting protein families. Protein families are often internally organized in distinct classes reflecting functional roles, domain architectures, phylogenetic relationships, or other biologically meaningful features. In bacterial two-component systems, for example, histidine kinases and response regulators can be grouped according to domain architecture, that is, the other domains present in the same polypeptide. Likewise, principal component analysis (PCA) projections of J-domain and Hsp70 sequences colored by InterPro-based architectural labels (Supplementary Figs. S1 and S2) and phylogenetic trees of the same families (Supplementary Fig. S3) reveal subdivisions into known subfamilies. When such labels are available, they provide a natural way to structure the matching problem.

In many cases, however, annotations are incomplete or unavailable, making it necessary to infer class membership directly from sequence information. We therefore evaluated several sequence-based classification strategies. The simplest, which is already reasonably effective, consists of applying PCA to one-hot encoded sequences and then clustering them with k-means. The number of clusters is selected by maximizing the silhouette score, which quantifies how well each sequence fits within its assigned cluster relative to neighboring clusters [49]. The corresponding silhouette scores and PCA projections for the datasets used here are shown in Supplementary Figs. S4 to S9. The class combinations observed among correct and incorrect pairs inferred from genomic proximity are shown in Supplementary Figs. S10 to S12.

Once classes have been assigned, CLAPP pools pairwise interaction scores across organisms at the class level through a pooling matrix (see section 5.2). Pooling aggregates information from many organisms, reducing fluctuations in the individual pairwise scores while also decreasing the dimensionality of the matching problem. The usefulness of this step relies on the assumption that class-level interaction patterns are sufficiently conserved across organisms to provide informative constraints on paralog matching.

In the present implementation, we restrict the sequences to be matched by CLAPP so that correct matches correspond to one-to-one interactions between classes. We therefore remove sequences belonging to classes that are represented multiple times within the same organism from the dataset of sequences to be matched. This one-to-one assumption, which also underlies most current paralog-matching algorithms, substantially simplifies the matching problem and allows us to isolate the contribution of class-level information. Importantly, this filtering applies only to the matching stage: sequences used to train the AttentionDCA model are not filtered in any way, and all possible within-species pairings are retained in the training set.

## 3 Results

### 3.1 Matching performance

We tested CLAPP on three bacterial datasets: histidine kinases (HK) and response regulators (RR), the ATPase and transmembrane components of the molybdate/tungstate ABC importer (ModBC), and J-domains with Hsp70s (see appendix S1.2). The ground truth for these datasets was assigned from genomic proximity.

We compared the performance of CLAPP against random matching (which reflects the difficulty of the dataset) and against the predictions of the Iterative Pairing Algorithm (IPA) [35], a common benchmark among paralog matching algorithms. Furthermore, we also report the prediction performance of the incomplete CLAPP, where the optimal class-to-class matrix is obtained organism-by-organism, without pooling. Fig. 2 shows matching performance on the tested datasets, with and without the pooling matrix implementation. CLAPP’s performance on additional families, also filtered so that correct matches correspond to one-to-one interactions between classes, is shown in Supplementary Fig. S13.

**Fig. 2:**
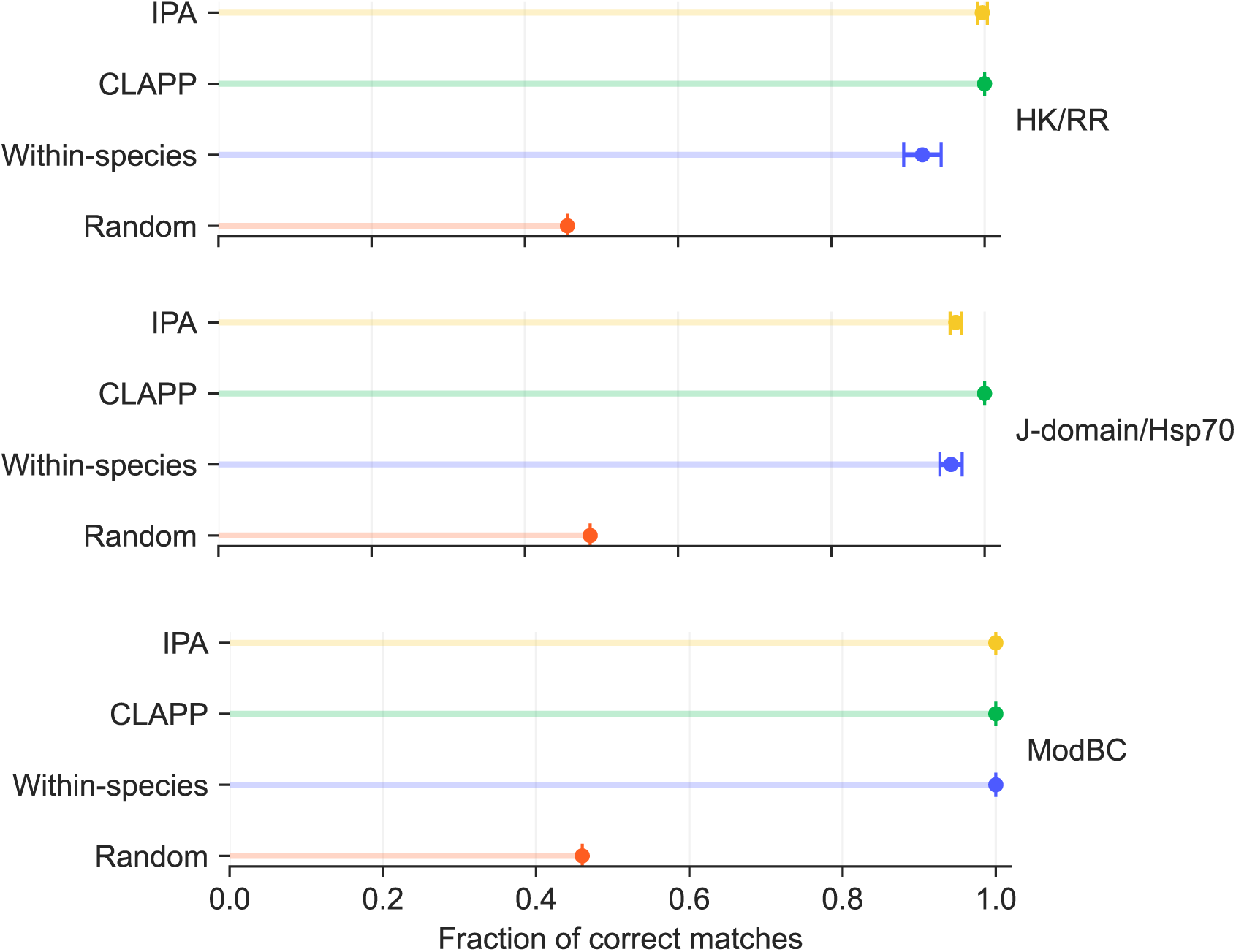
Paralog matching performance of AttentionDCA scores within-species and with pooling matrix (CLAPP). Sequences are divided into clusters using k-means and cluster labels are used for the pooling matrix. Performance is compared to the iterative pairing algorithm (IPA) [35] and to the random assignment of within-species pairs. Results are averaged over five random initializations, with error bars indicating one standard deviation.

Across all datasets, pairwise scores performed above random, but perfect matching was achieved only after incorporating sequence-class information through pooling. CLAPP thus turns imperfect pairwise evidence into exact assignments by exploiting the collective organization of protein families across organisms. At the same time, this highlights that CLAPP requires sufficiently accurate and unambiguous class assignments; errors or uncertainty in these labels can affect matching performance.

### 3.2 Sensitivity of CLAPP to Class-Label Quality

To assess CLAPP’s sensitivity to class definition, we evaluated it on an unfiltered Hsp70/J-domain dataset containing all sequences for which genomic proximity could be used as a ground truth for interactions. Unlike the filtered datasets used above, sequences were not selected to enforce one-toone interactions between classes, nor were organisms containing multiple representatives of the same class excluded.

For this unfiltered dataset, class labels were initially assigned using PCA followed by k-means clustering. We then generated a second version of the same dataset in which these labels were refined using phylogenetic information. Separate phylogenetic trees were constructed for Hsp70s and J-domains, including reference sequences with known subfamily labels (DnaK, HscA and HscC for Hsp70s; DnaJ, HscB and DjlC for J-domain proteins; see appendix S1.2) together with the unfiltered sequences to be paired. The resulting trees are shown in Supplementary Fig. S3.

We compared the k-means assignment of each sequence with its position in the corresponding phylogenetic tree. Whenever a sequence grouped within a phylogenetic clade inconsistent with its k-means label, we reassigned it accordingly. For example, in the Hsp70 tree several sequences assigned to cluster 4 grouped with HscA sequences and were therefore reassigned to cluster 1, which is associated with HscA in both the phylogenetic tree and the PCA projection.

This procedure produced a curated dataset in which class labels reflect both sequence-space proximity and evolutionary relationships. After relabeling, the distribution of correct J-domain/Hsp70 pairs across class combinations became substantially cleaner (Supplementary Fig. S14), consistent with the expectation that interactions between classes are conserved across organisms.

Both the PCA projections (Supplementary Fig. S15) and the Hsp70 phylogenetic tree (Supplementary Fig. S3) indicate that Hsp70 k-means clusters 2 and 3 correspond to DnaK sequences. The interaction data agree with this assignment: both clusters interact exclusively with J-domain cluster 3, corresponding to DnaJ. We therefore merged these two Hsp70 classes before applying the pooling procedure.

Fig. 3 compares CLAPP performance using the original k-means labels and the phylogeny-informed labels. Performance with the k-means labels is substantially lower than on the filtered dataset (Fig. 2, middle panel). This reflects two sources of difficulty. First, the unfiltered dataset violates the assumptions underlying the pooling procedure: interactions between classes are not necessarily one-to-one, and individual organisms may contain multiple representatives of the same class. Indeed, the presence of multiple representatives of the same class within an organism introduces an additional ambiguity that cannot be resolved from the class-to-class interaction map alone. Second, an incorrectly labeled sequence can place a correct pair in a class combination inconsistent with the dominant class-level interaction pattern, causing the pooled assignment to disfavor it.

**Fig. 3:**
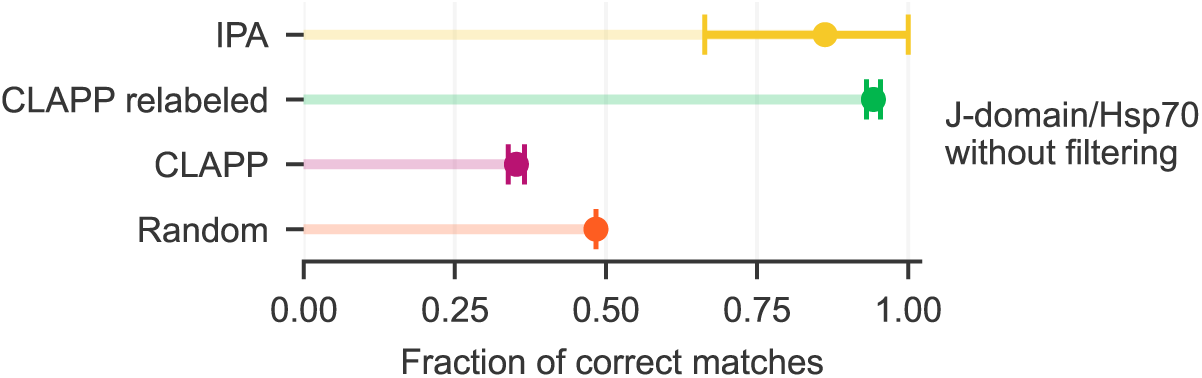
Paralog matching performance of AttentionDCA scores on bacterial Hsp70s and J-domains with pooling matrix (CLAPP). The methods are tested on 320 pairs of interacting sequences. Sequences are divided into clusters using k-means (”CLAPP” results) and relabeled if needed following their position in the family phylogenetic tree (”CLAPP relabeled” results). Cluster labels are used for the pooling matrix. Performance is compared to the iterative pairing algorithm (IPA) [35] and to the random assignment of within-species pairs. Results are averaged over five random initializations, with error bars indicating one standard deviation.

Phylogeny-informed relabeling markedly improves performance. On the relabeled dataset, CLAPP correctly predicts on average 94.3% of the pairs inferred from genomic proximity. These results show that CLAPP can remain highly accurate on substantially more complex datasets, but in that case the quality of the class labels becomes critical.

### 3.3 Residual Errors Reflect Within-Class Interaction Ambiguity

After phylogeny-informed relabeling, all remaining incorrect CLAPP assignments involve classes corresponding to DnaK and DnaJ. In every affected organism, multiple representatives of these classes are present, and CLAPP selects DnaK-DnaJ pairings that are not supported by genomic proximity.

These residual errors are therefore qualitatively different from those caused by noisy class labels. In these cases, the assumption of a unique interaction partner may itself be questionable, because DnaK and its JDP co-chaperones can interact promiscuously. One relevant example is CbpA, a JDP whose function partially overlaps with DnaJ [50] and which our labeling procedure classifies with DnaJ. The fraction of organisms in which CLAPP makes at least one incorrect assignment is 0.32 among organisms containing at least one CbpA, compared with 0.01 among organisms without CbpA. This enrichment suggests that at least some disagreements with genomic proximity occur in systems where non-exclusive interactions are biologically plausible.

A second indication comes from sequence similarity. For each protein, we compare its sequence with those of the other paralogs of the same family in the same organism. Proteins from pairs that CLAPP predicts correctly across all random initializations are, on average, more distant from these alternative paralogs than proteins from misassigned pairs. For J-domains from correctly assigned pairs, the mean Hamming distance to alternative paralogs is 0.72, compared with 0.43 for J-domains from misassigned pairs. Similarly, Hsp70s from correctly assigned pairs have a mean Hamming distance of 0.60 to alternative paralogs, compared with 0.41 for Hsp70s from misassigned pairs. Misassignments are therefore concentrated among closely related paralogs, consistent with incomplete functional specialization.

IPA [35] shows a different failure pattern on this dataset. Its performance is substantially more sensitive to random initialization, with a core set of consistently misassigned pairs plus additional initialization-dependent errors. These errors involve proteins from all classes, and we did not identify a common structural pattern underlying them. Moreover, they do not coincide with the pairs misassigned by CLAPP.

Taken together, these observations suggest that, once class labels are reliable, the residual errors of CLAPP are concentrated in cases where the underlying biological interaction map may not be strictly one-to-one. Such cases represent an inherent limitation of the assumption that interactions between paralogs are one-to-one, shared by most paralog matching methods.

### 3.4 Predicting eukaryotic complex structures using CLAPP

In appendix S1.1, we show that CLAPP remains useful beyond highly curated benchmarks: in these unfiltered eukaryotic datasets, CLAPP-based pairing performed comparably to standard AlphaFold3 pairing. This suggests that the method is sufficiently robust to provide informative pairings even when its assumptions are only approximately satisfied.

### 3.5 Emergence of the correct assignment

The striking performance of CLAPP, which achieves perfect matching across the curated datasets and remains highly accurate even in the presence of biologically plausible promiscuity, motivates a closer examination of how the correct assignment emerges. To understand how the relevant signal is extracted from the AttentionDCA interaction energies, we performed additional analyses varying both the training hyperparameters and the composition of the input MSA. These tests address two questions: how rapidly the correct class-level assignment appears during training, and how much information about the true pairings is required for this transition to occur.

#### 3.5.1 Early emergence of the correct class assignment

Fig. 4 shows CLAPP’s matching performance on the HK/RR and J-domain/Hsp70 datasets as a function of the number of AttentionDCA training epochs. During each epoch, the entire training dataset is processed once in batches, with model parameters updated by gradient-based optimization after each batch. Strikingly, a single epoch is sufficient for CLAPP to reach 100% accuracy for both datasets, indicating that the information required for class-level partner assignment emerges extremely early during training.

**Fig. 4:**
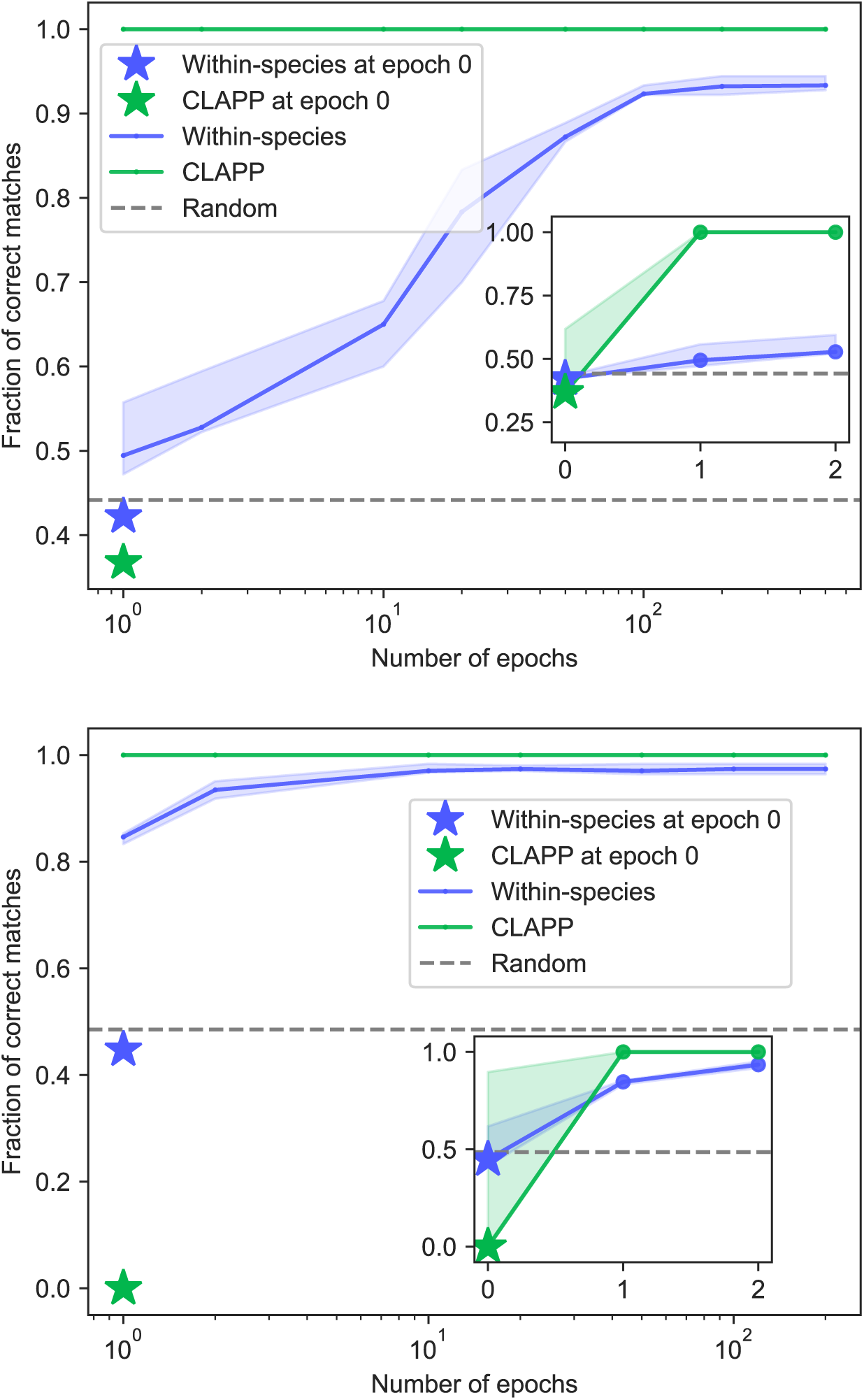
Paralog matching performance as a function of AttentionDCA training epochs. Fraction of correct matches obtained using AttentionDCA scores for the HK/RR dataset (top) and bacterial J-domain/Hsp70 dataset (bottom), either by independent within-species matching or after class-level pooling with CLAPP. Epoch 0 corresponds to the randomly initialized AttentionDCA model, before any parameter update. Insets show performance at 0,1, and 2 epochs, while main panels show performance starting from 1 epoch in log scale, with performance at epoch 0 also shown as a reference. Sequences are divided into classes using k-means, and class labels are used to construct the pooling matrix. Results are shown for five random initializations; solid lines indicate the median and shaded regions the interquartile range.

We repeated the analysis for different learning rates (Supplementary Fig. S16). In the absence of parameter updates, i.e. for vanishing learning rates, CLAPP remains close to random performance. For the smallest non-zero learning rate tested, the correct assignment emerges only after several epochs, whereas larger learning rates recover it after the first epoch. Thus, the transition appears to occur once the cumulative parameter updates are sufficient to reorganize the class-level cost matrix so that the correct assignment becomes optimal.

To characterize this reorganization more directly, we considered the pooled cost matrix *C* and examined how strongly it favors the optimal assignment over alternative ones. Within CLAPP, the class-level assignment is represented by a permutation matrix *S*, and the optimal assignment minimizes

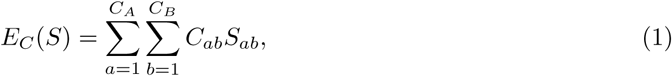

with *C_A_*and *C_B_*the number of classes in families A and B respectively. We associate to each possible permutation a Boltzmann-like probability

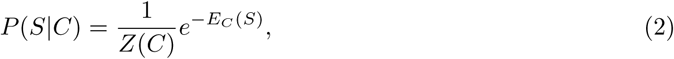

with

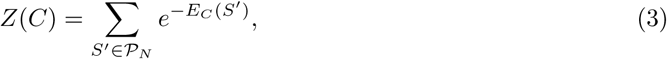

where the sum runs over all permutation matrices of size *N* , the number of classes. We then define

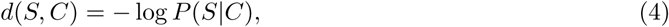

which measures how strongly a given cost matrix disfavors a permutation: smaller values of *d*(*S, C*) correspond to assignments that are more strongly favored by *C*. This quantity therefore reports not only which assignment is optimal, but also how well it is separated from competing permutations.

The left panel of Fig. 5 shows *d*(*S, C*) for all possible permutations at different training epochs for the HK/RR dataset. A clear qualitative change occurs between epochs 0 and 1. At initialization, the different permutations have similar values of *d*(*S, C*), whereas after a single epoch the correct permutation becomes the minimum and is clearly separated from the alternatives. With further training, the correct permutation remains optimal, but its separation from competing assignments progressively decreases before approaching a stationary profile. At the same time, variability across random initializations decreases, indicating that prolonged training makes the resulting assignment landscape more reproducible. The right panel of Fig. 5 shows the fraction of correctly predicted contacts by AttentionDCA models trained for increasing numbers of epochs on the HK/RR dataset. The fraction sharply increases in the initial epochs and reaches a plateau around epoch 50.

**Fig. 5:**
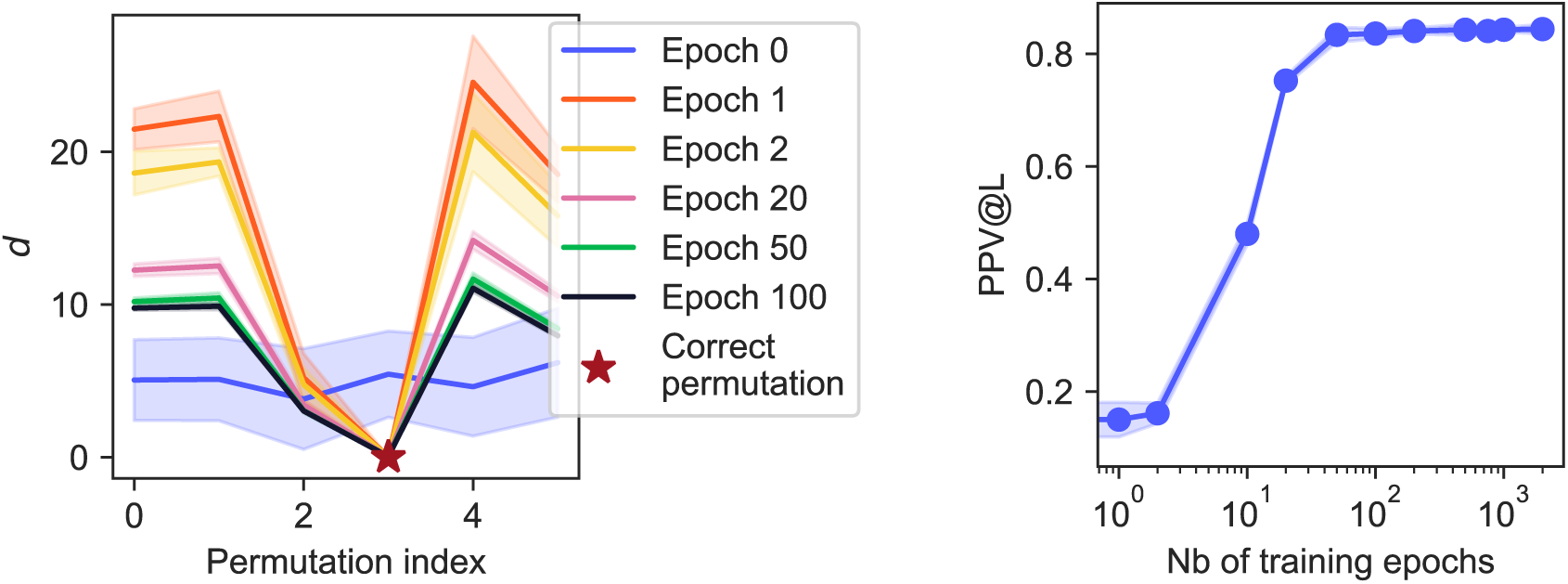
Evolution of class-assignment and contact-prediction signals during AttentionDCA training. Left panel: *d*(*S, C*) = *−* log *P* (*S | C*) for each possible class-level permutation *S*, computed from pooled cost matrices *C* obtained after different numbers of AttentionDCA training epochs on the HK/RR dataset. Lower values indicate permutations that are more strongly favored by the cost matrix. Right panel: positive predictive value (PPV), defined as the fraction of true contacts among the top *L* predicted contacts, where *L* is the length of the MSA, as a function of the number of training epochs. Solid lines and shaded regions show the mean and standard deviation, respectively, across five random initializations.

The same behavior can be summarized through the statistical free energy

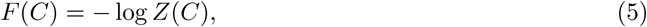

and its difference from the minimum assignment cost, *F* (*C*) *− E*^min^. Writing *S*^opt^ for the optimal permutation,

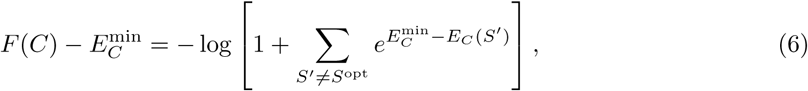

so that this quantity reflects the statistical weight carried by assignments competing with the optimum.

As shown in Fig. 6, *F* (*C*) reaches a minimum after one training epoch and subsequently increases toward a plateau. Similarly, *F* (*C*) *− E*^min^ is closest to zero after one epoch, indicating that the optimal assignment carries its largest relative weight at this point. Subsequent training preserves the correct assignment but increases the relative weight of competing permutations.

**Fig. 6:**
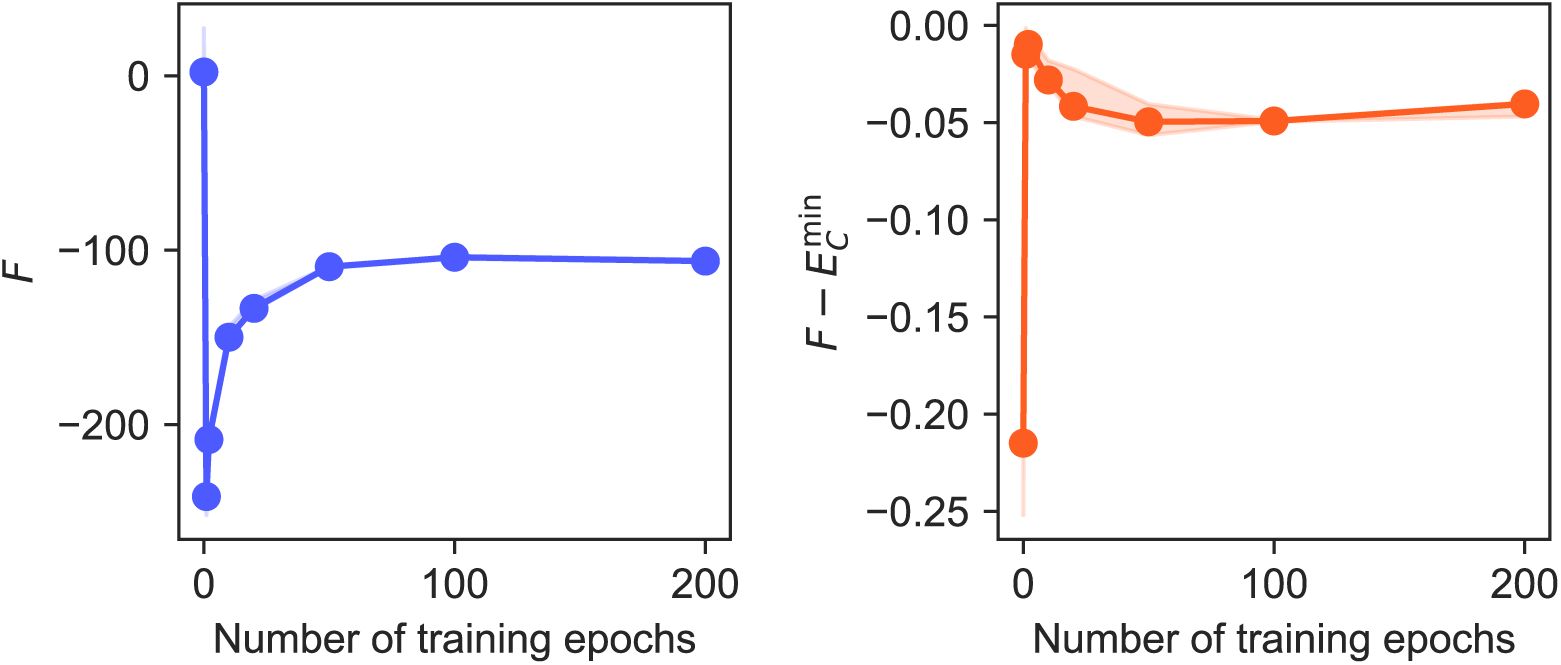
Evolution of the class-assignment free-energy landscape during AttentionDCA training. Statistical free energy *F* (*C*) (left) and contribution of non-optimal assignments, quantified by *F* (*C*) *− E*^min^ (right), for pooled cost matrices obtained after different numbers of AttentionDCA training epochs on the HK/RR dataset. Results are shown for five random initializations; solid lines indicate the median and shaded regions the interquartile range.

Taken together, these results reveal a separation between the training timescales of class-level partner assignment and residue-level contact prediction. The signal required to identify the correct class assignment is already present after a single epoch, whereas contact-prediction performance continues to improve until approximately epoch 50. Further training reduces variability across random initializations, making the assignment landscape more reproducible, but does not improve the assignment itself and progressively reduces the separation between the correct permutation and its competitors. This behavior is consistent with AttentionDCA learning coarse class-level interaction preferences before refining residue-level couplings.

#### 3.5.2 How much pairing information is required?

Having established that the correct class assignment emerges very early during training, we next asked how much information about the correct interacting pairs is required for this transition to occur.

Fig. 7 (left panel) and Supplementary Figs. S17 to S19 show matching performance for AttentionDCA models trained on MSAs of increasing depth for the HK/RR and J-domain/Hsp70 datasets. Performance improves rapidly with the number of sequences used for training, and CLAPP reaches 100% accuracy with surprisingly small training MSAs. This contrasts with coevolutionary models used for residue-contact prediction, which generally require substantially deeper alignments [17, 51, 52].

**Fig. 7:**
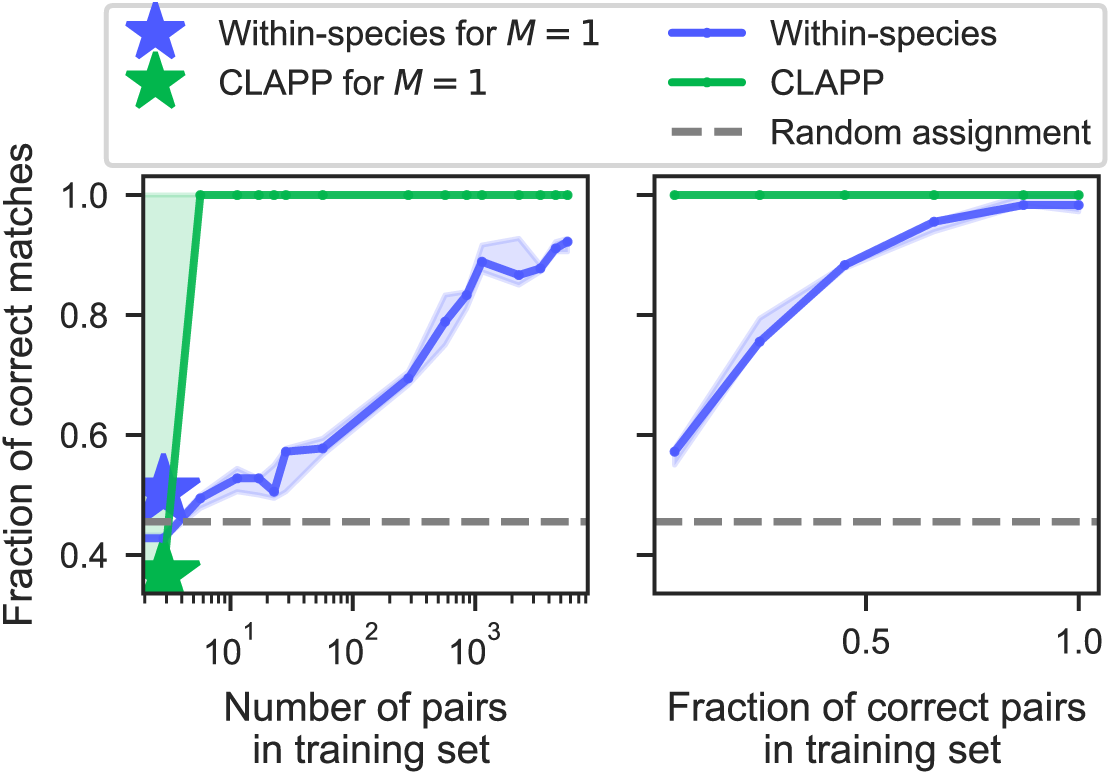
Paralog matching performance as a function of training MSA size (left) and fraction of correct pairs in the training MSA (right). Fraction of correct matches obtained using AttentionDCA scores for the HK/RR dataset, either by independent within-species matching or after class-level pooling with CLAPP. Left panel: for each training set size, a subset of interacting pairs was selected and all possible within-species pairings among the corresponding sequences were included in the AttentionDCA training MSA. The panel shows performance starting from 1 pair in log scale, with performance for 1 sequence in the training set also shown as a reference. Right panel: the training MSAs for AttentionDCA are kept at fixed total depth, while varying the fraction of correct pairs in them. Sequences are divided into classes using k-means, and class labels are used to construct the pooling matrix. Results are shown for five random initializations; solid lines indicate the median and shaded regions the interquartile range.

We then fixed the total depth of the training MSA while progressively reducing the fraction of correct pairs. As shown in the right panel of Fig. 7, CLAPP remains highly accurate even as the number of correct pairs decreases. Remarkably, an MSA containing no correct pairs can still produce a model from which CLAPP recovers the correct class-level assignment. Supplementary Fig. S20 shows the corresponding results when AttentionDCA is trained exclusively on incorrect pairs for zero or one epoch. After a single training epoch, four out of five random initializations recover the correct assignment with 100% accuracy.

To understand this apparently counterintuitive result, we examined the role of the prior used when constructing the pooled class-level matrix. As described in section 5.2, each candidate pair ***a^m^*** is weighted by

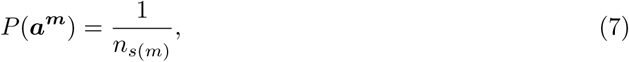

where *n_s_*_(*m*)_ is the number of possible paralog partners in organism *s*(*m*). This prior gives greater weight to candidate pairs from organisms in which the matching problem contains fewer alternatives. Fig. 8 compares the assignment landscapes obtained from AttentionDCA models trained exclusively on correct or incorrect pairs, with and without this prior. When the model is trained on correct pairs, *d*(*S, C*) displays a pronounced minimum at the correct class permutation and the overall landscape is essentially unaffected by the prior. In this regime, the interaction score itself already contains a strong signal for the correct assignment.

**Fig. 8:**
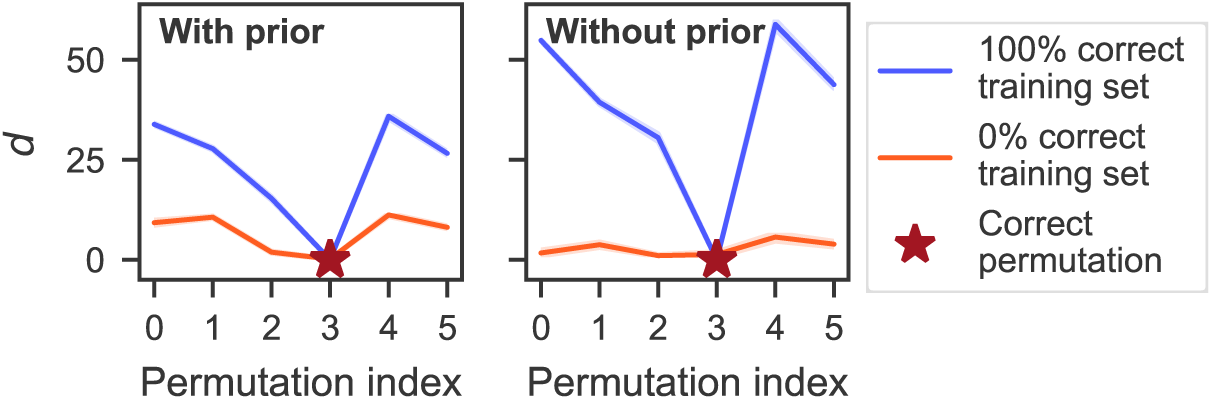
Effect of the pairing prior on the class-assignment landscape. Values of *d*(*S, C*) = *—* log *P* (*S|C*) for all possible class-level permutations *S*, using pooled cost matrices *C* obtained from AttentionDCA models trained exclusively on either correct pairs or incorrect pairs, with (left panel) and without (right panel) the prior on pair correctness. Lower values indicate permutations that are more strongly favored by the cost matrix. Results are shown for five random initializations; solid lines indicate the median and shaded regions the interquartile range.

The situation is different when AttentionDCA is trained exclusively on incorrect pairs. In this case, the correct class assignment is recovered only when the prior is included. The prior therefore has little influence when the sequence-derived interaction score is already informative, but becomes important when that signal is weak or poorly learned. By weighting more strongly organisms with fewer possible partners, it allows the pooling procedure to extract a consistent class-level signal even from a model trained without explicit examples of the correct pairs.

These results show that CLAPP requires surprisingly little direct information about the correct pairings in the training MSA. When the interaction model already captures the relevant signal, the prior is largely dispensable; when that signal is weak, the combination of class-level pooling and the organism-level pairing prior can nevertheless recover the correct interaction pattern.

#### 3.5.3 What sequence information is required?

The results above show that CLAPP can recover the correct class-level assignment very early during AttentionDCA training and even when the training MSA contains little or no information about the correct pairings. This raises a more fundamental question: which sequence features are actually required for successful matching?

To address this question, we replaced the AttentionDCA interaction energy with scores derived from two progressively simpler models (see section 5.1.2). The first, which we call the *fields-only* model, retains only site-specific amino-acid conservation and contains no couplings between sequence positions. The second, which we call the *composition* model, discards positional information altogether and compares sequences only through their amino-acid compositions.

Supplementary Fig. S21 compares the matching performance of these scores with that obtained from AttentionDCA across the three bacterial datasets considered in section 3.1. Strikingly, the fields-only score is sufficient for CLAPP to recover the correct class-level interaction pattern, and consequently the correct partner assignments, in all three datasets. The composition score is less robust: it performs well on the HK/RR and ModBC datasets, but fails to recover the correct assignment for the J-domain/Hsp70 system.

The success of the fields-only model is particularly notable because it contains no explicit coevolutionary information between the two protein families. Moreover, when the same score is used to perform matching independently within each organism, its performance is poor (Supplementary Fig. S21). Thus, site-specific conservation alone does not generally identify the correct individual partners. Rather, its signal becomes informative when aggregated across organisms according to the class structure.

As a further control, we considered the fields extracted from the AttentionDCA couplings during the gauge transformation used to calculate the interaction score. This transformation separates the learned couplings into effective single-site fields and residual gauge-fixed couplings, with the standard AttentionDCA score retaining only the latter. Remarkably, a score constructed using only the extracted fields also allowed CLAPP to recover the correct matches. Thus, the matching information needed to correctly identify interaction partners is present in both the effective one-body component extracted by the gauge transformation and in the residual coupling component obtained after the gauge, either of which is sufficient for successful matching after class-level pooling.

These results further highlight the central role of pooling in CLAPP. Explicit inter-protein coevolutionary couplings are not always required to recover the correct class-level interaction map: sufficiently informative sequence features, even considerably simpler ones, can become predictive once evidence is combined across organisms. At the same time, the failure of the composition score on the J-domain/Hsp70 dataset shows that this information cannot always be reduced to very coarse sequence properties, and that the level of sequence detail required depends on the protein families considered.

## 4 Discussion

In this work we introduced CLAPP, a method for paralog matching that leverages conserved interactions between subclasses within interacting protein families. Rather than solving a separate one-to-one assignment problem for each species, CLAPP aggregates interaction scores across organisms for pairs of classes and infers a single class-to-class interaction matrix. This reduces sensitivity to noisy scores and replaces an organism-level permutation problem with a much smaller assignment at the class level. Across the systems tested, class pooling leads to perfect recovery of the ground-truth assignments.

In families with multiple paralogs per genome, assigning interaction partners is necessary to understand what drives interactions and to better study how pairs evolve to play different functions. In addition, accurate partner assignment is required to curate high-quality paired datasets for downstream modeling: both explicit coevolutionary models and deep learning models benefit from correct pairs for tasks such as complex structure determination or design. From this perspective, CLAPP is also a first step for building reliable training resources and for interpreting how specificity is encoded in sequences.

Sequences can be classified using architectural or functional labels derived experimentally. When these labels are not available, in the current pipeline, sequences are clustered using PCA followed by k-means and further filtered to retain cases consistent with a one-to-one mapping between classes and to exclude organisms containing multiple sequences from the same class. This filtering facilitates the perfect performance of CLAPP by enforcing the core assumption behind it: conserved, exclusive class-level interactions. However, the pooling framework itself does not intrinsically require organisms to only have one representative per class.

While using class information improves paralog matching, the current implementation of CLAPP has two main limitations. First, its performance depends on the quality of the class assignments and on the existence of biologically meaningful subclasses with conserved interaction patterns across organisms. Although unsupervised clustering can recover known subtypes, as shown for J-domains, misclassification can worsen pooling quality. This is illustrated by the improved matching performance obtained after phylogeny-informed relabeling of the Hsp70/J-domain dataset, showing that better class assignments translate directly into better matching.

Second, the current method assumes one-to-one interactions between classes. Experimental evidence, however, shows that proteins can have multiple partners, resulting in promiscuous interactions and crosstalk that violate this assumption at either the paralog or class level [12, 53].

Although the reliance on class labels is specific to CLAPP, it reflects a broader challenge shared by paralog-matching methods, namely sensitivity to the quality of the signals used to guide partner assignment. By contrast, the assumption of exclusive one-to-one interactions is a limitation shared explicitly by most current approaches. In fact, several apparent CLAPP errors occur in systems showing evidence of interaction promiscuity, suggesting that they may partly reflect a mismatch between this constraint and the underlying biology. Extending the framework to accommodate non-exclusive interactions could therefore produce assignments that better capture biological interaction networks.

While investigating how much information CLAPP needs to correctly identify interaction partners, we found that explicitly coevolution-aware models are not needed to achieve good performance. In fact, a fields-only or profile model is sufficient to correctly identify interactions between classes. This shows the power of the pooling mechanism underlying CLAPP, which allows correct pair recovery even when interaction scores are calculated from extremely simple models.

Overall, CLAPP provides a computationally efficient and score-agnostic mechanism to exploit conserved subclass structure for paralog matching. When class-level interaction organization is present, pooling can dramatically improve pairing accuracy and reduce computational cost. Beyond improving matching itself, the method can facilitate the construction of correctly paired datasets needed to study the determinants of interaction specificity and to enable sequence-based prediction and design of interacting protein partners.

## 5 Methods

### 5.1 Interaction scores

#### 5.1.1 Factored attention

We use an interaction score derived from Direct Coupling Analysis (DCA), which defines a statistical energy over a multiple sequence alignment (MSA). Specifically, we train a factored-attention DCA model (AttentionDCA) [45, 46] on a concatenated alignment of proteins from interacting families A and B, constructed from all possible within-species A–B sequence concatenations, including both correct and incorrect pairings. This strategy makes it possible to infer interacting partners without requiring a reference set of known pairs, and can therefore be applied even to families for which interaction assignments are unavailable.

From the couplings learned by the model, to which the zero-sum gauge is applied, we define the interaction energy [35] of any concatenated sequence pair as

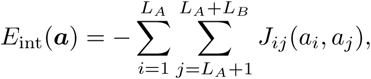

where ***a*** = (*a*_1_*, . . . , a_L__A_ , a_L__A_*_+1_*, . . . , a_L__A_*_+*L*_*_B_* ) denotes a concatenated pair of aligned sequences from families A and B, with *a_i_, a_j_ ∈ {*1*, . . . , q}*, *q* = 21, and where *L_A_* and *L_B_* are the lengths of the aligned sequences from families A and B, respectively.

In the factored-attention DCA model, the Potts couplings are parametrized as

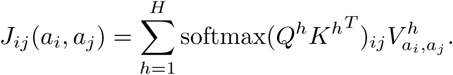

Compared with more traditional implementations of Potts-model DCA, this parametrization offers several advantages. In particular, the low-rank structure of the couplings substantially reduces the number of parameters while retaining strong performance on tasks such as contact prediction [46]. In the present context, AttentionDCA also provides a flexible framework that could naturally be extended to learn the pooling matrix during training, opening the way to an end-to-end differentiable inference of interaction partners.

When using the negative AttentionDCA interaction energy as the interaction score, we set the number of heads to *H* = 150, the internal dimension to *d* = 23, and determine the number of training epochs by early stopping.

#### 5.1.2 Profile and composition models

Other interaction scores tested rely on simpler models that do not take into account coevolution. First, we considered a score constructed from the fields of a profile model *H*(***a***) = *−* ^Σ^*_i_ h_i_*(*a_i_*), with fields calculated from the empirical frequencies of the MSA *f_i_* using a pseudocount correction [16] *f̄_i_*(*a*) = (1 *−* Λ)*f_i_*(*a*) + Λ*/q*, Λ = 0.5: *h_i_*(*a*) = ln *f̄_i_*(*a*). This model captures site-specific conservation but contains no information about correlations between positions and therefore cannot describe coevolution. From the fields, we define the interaction energy

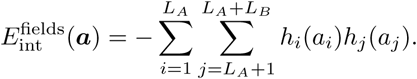

where ***a*** = (*a*_1_*, . . . , a_L__A_ , a_L__A_*_+1_*, . . . , a_L__A_*_+*L*_*_B_* ) denotes a concatenated pair of aligned sequences from families A and B, with *a_i_, a_j_ ∈ {*1*, . . . , q}*, and where *L_A_* and *L_B_* are the lengths of the aligned sequences from families A and B, respectively. Terms in this score are equivalent to the couplings in the Hopfield-Potts model [54] with a single attractive pattern.

Following the rearrangement inequality, the total interaction energy over a one-to-one assignment is minimized by pairing sequences with similarly ranked profile scores, i.e. pairing the A sequence with highest value of 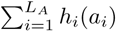 with the B sequence with highest 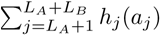, pairing the ones with second highest values, etc. Since the sum of fields is a measure of how closely each sequence conforms to the position-specific frequencies of the training MSA, the model matches sequences according to their profile typicality. Since this may reflect shared phylogenetic divergence, this score can act as a proxy for phylogenetic information.

Looking for an even simpler model, we defined one where also the site information is lost and only sequence composition (*i.e.* which amino acids are present in each sequence, irrespective of their position in the sequence) is taken into account. In particular, we define 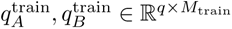 the matrices containing the amino acid frequencies for each training sequence. Then 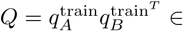 R*^q^*^×^*^q^* is a compatibility matrix of amino acid compositions, with *Q_ab_* indicating whether amino acid *a* in a sequence from family A is compatible with amino acid *b* in a sequence from family B. Finally, we take

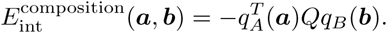

### 5.2 Pooling matrix

Proteins within a family can often be partitioned into subclasses or functional groups. If interactions between classes are conserved across organisms, *i.e.* if sequences belonging to one class only interact with sequences from another class, then this information can be used to simplify the paralog matching problem. Rather than inferring interactions directly at the level of individual paralog pairs, we introduce a class-level coupling matrix *S ∈* R*^CA^*^×^*^CB^* , where *C_A_* and *C_B_* denote the numbers of classes in families A and B, respectively. Under the assumption that sequences belonging to the same class share similar interaction preferences, paralog matching can then be reformulated in terms of inferring the interactions between classes. This replaces a sequence-level assignment problem over *M* candidate pairs by a much smaller class-level optimization problem, thereby substantially reducing the computational cost.

For a set of candidate pairs (***a****^m^*)_m=1_*^M^* , where each pair ***a****^m^*is formed by one sequence from family A and one from family B, we define the class-level score matrix *C ∈* R*^CA^*^×*CB*^ as

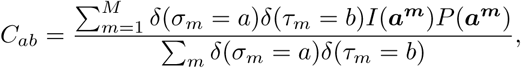

where *σ_m_* and *τ_m_* are the class labels of the A and B sequences in pair *m* and *I*(***a****^m^*) is the interaction score of pair ***a****^m^*. We set

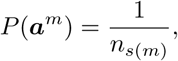

where *n_s_*_(*m*)_ is the number of possible paralog partners in species *s*(*m*). This factor represents the prior probability that a candidate pair is correct under a uniform prior over the possible within-species partners. The matrix *C* can therefore be interpreted as an estimate of the propensity of class *a* in family A to interact with class *b* in family B, obtained by averaging pairwise interaction scores over all candidate pairs assigned to that pair of classes.

Our goal is then to infer a matrix *S* that best captures the class-to-class interaction structure by maximizing

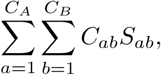

subject to the marginal constraints

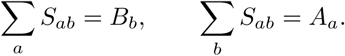

This optimization problem can be solved exactly and in most cases efficiently by linear programming [47, 48]. When *C_A_* = *C_B_*, choosing uniform marginals *A_a_* = 1*/C_A_* for all *a* and *B_b_* = 1*/C_B_* for all *b* yields the feasible set of scaled doubly stochastic matrices, whose optimal solution corresponds to a scaled permutation matrix in the absence of degeneracies.

### 5.3 Comparison with the iterative pairing algorithm

We compare CLAPP to the iterative pairing algorithm (IPA) based on the mean-field approximation [35]. IPA infers, within each organism, one-to-one matches represented by permutation matrices and iteratively constructs DCA models from the corresponding concatenated alignments. For each pair of interacting families, the input alignments include both the sequences used to train AttentionDCA and those that were paired using CLAPP, while performance is evaluated on the sequences on which CLAPP was tested only. When the numbers of sequences in the two families differ within a training organism, the smaller family is padded with gap-only sequences.

### 5.4 Phylogenetic analysis of selected proteins

Phylogenetic analyses of Hsp70s (i.e. DnaK, HscA, HscC) homologs and corresponding J-domains of JDPs (i.e. DnaJ, HscB, DjlC) were performed using IQ-TREE 3.1.0 [55]. The protein sequences were aligned using the methodology described under the relevant section. The best-fitting substitution model was determined by ModelFinder [56] as Q.PFAM+I+G4, which was subsequently used for maximum likelihood tree reconstruction. Branch support was estimated using 1,000 ultrafast bootstrap replicates [57] and approximate likelihood-ratio test (aLRT) [58]. The resulting trees were visualized and annotated in iTOL [59].

## S1 Supplementary Materials

### S1.1 Eukaryotic Complex Structure Prediction

Predicting eukaryotic protein-complex structures remains particularly challenging because constructing informative paired MSAs, which are required by most state-of-the-art methods, is often difficult. Eukaryotic protein families typically contain more paralogs per species than their prokaryotic counterparts, substantially increasing the number of possible pairings. At the same time, fewer homologous sequences are generally available for eukaryotic protein families, and their alignments tend to exhibit lower sequence diversity, providing less information from which to infer inter-protein coevolution [39, 60]. Moreover, unlike many prokaryotic interaction partners, eukaryotic partners are generally not encoded in conserved operons, so genomic proximity cannot be used as a proxy for the correct pairing. Incorrectly paired sequences in concatenated MSAs dilute the inter-protein coevolutionary signal supplied to structure-prediction models, potentially preventing accurate reconstruction of the interaction interface. Improving paralog matching is therefore an important step towards extending reliable structure prediction to eukaryotic complexes and elucidating the molecular mechanisms of these assemblies.

To test whether our paralog matching methods can improve eukaryotic heterodimer structure prediction, we selected structures from the Protein Data Bank (PDB) from the dataset curated by [61] of structures released after the AlphaFold 3 (AF3) [62] training cutoff dates and for which AF3 does not predict correct structures. In practice, this was done by selecting structures for which the model had reported ipTM scores below 0.5, as ipTM is a good proxy for model accuracy [61]. An additional selection was performed to retain only heterodimers formed by two distinct proteins from the same eukaryotic species. Complexes were excluded when one chain originated from a different species, when the interaction clearly depended on a non-physiological ligand not present in cells, or when the two chains corresponded to the same protein. Structures were then predicted using a modified version of the AF3 ColabFold notebook [15, 62], with and without an MSA paired using CLAPP. When ligands or ions were present in the PDB, they were also supplied to the model. Supplementary Tab. S1 summarizes the selected PDB entries and the characteristics of the corresponding CLAPP-paired MSAs.

For each structure, we retrieve sequences belonging to each of the protein families forming the complex using MMseqs2 [63, 64]. In the ColabFold pipeline, sequences in each species and family are sorted by increasing E-value, and the pair with highest scores in each organism is added to the MSA used as input to the model. We paired sequences in each organism using CLAPP and added sequences made of only gaps when the two families contained a different number of sequences in a given organism. The AttentionDCA model was trained on all possible within-organism matches. We did not match sequences from organisms with more than 30 sequences in either family. The resulting MSA is then used as input for AF3. Five samples are generated for each settings configuration with ten recycles and no templates. Supplementary Fig. S22 compares complex structures predicted with and without a CLAPP-paired MSA, as well as a randomly generated paired MSA, based on local model confidence (mean pLDDT [65]), confidence in the relative placement of the interacting chains (ipTM [66]), and agreement with the experimental structure (DockQ [67, 68]).

For most targets, replacing the standard MSA pairing with CLAPP pairing yields only modest improvements, if any, in complex-structure prediction. This is particularly evident from the DockQ scores, which quantify the agreement between the predicted and experimental interfaces [67, 68]. Some predictions (7ZHR, 8BZR and 8HLN) obtain low DockQ scores, indicating that the experimental binding mode is not accurately recovered. These results suggest that paralog matching may not be the main bottleneck for these targets. Prediction accuracy may instead be limited by insufficient sequence depth or diversity or weak evolutionary constraints at the interface. A further possibility is that the experimental structure captures only one of several accessible conformational or interaction states, with the prediction method favouring another [69, 70]. In such cases, a low DockQ score may reflect a difference between the predicted and experimentally observed states rather than an intrinsically implausible interface. More generally, evaluating complex predictions against a single experimental reference can be problematic when heterodimers adopt multiple configurations [61,71]. In other cases, such as 7S7L and 7VVB, supplying AF3 with the ligands present in the experimental PDB substantially improves the prediction scores, indicating that these ligands play an important role in stabilizing the experimentally observed complex structure. CLAPP’s performance for eukaryotic complex structure prediction is compatible with DiffPALM’s performance [39], which did not improve scores when compared to the ColabFold default pairing method.

### S1.2 Datasets

The HK–RR and ModBC datasets were assembled by assigning candidate interacting pairs from genomic proximity [35,36,72–74], whereas the J-domain/Hsp70 dataset was assembled here, also from genomic proximity. For this dataset, only the J-domain region of JDPs was retained in the alignments and used throughout the analysis, since the relevant phylogenetic and functional information is known to be encoded within the J-domain itself [75].

To train the AttentionDCA model, we use alignments constructed from all possible within-organism sequence pairs. This makes it possible to train the model without prior knowledge of which paralogs interact, relying instead on the fact that the correct pairs are contained among the full set of candidate pairs. For this strategy to be valid, the relevant paralogs from both families must be represented as completely as possible in each organism.

To this end, we created a database of bacterial proteomes from UniProt with BUSCO score *>* 90 (to ensure the completeness of a proteome assembly) [76–78] and standard Complete Proteome Detector score [79].

For the Hsp70 and J-domain families, the training set was constructed by selecting all sequences in this curated proteome database annotated with InterPro identifiers IPR013126 and IPR036869, respectively. These sequences were then aligned and used to generate all possible within-organism A-B concatenations for model training. In total, training was done using 4287 Hsp70 sequences and 7428 J-domain sequences. For HK/RR, we used 5680 sequence pairs and for ModBC 4132.

For evaluation of Hsp70/J-domain pairs, we used bacterial reference proteomes with annotated genomic positions. Sequences were again selected using the same InterPro identifiers, and likely interacting pairs were inferred from genomic proximity, with pairs separated by more than ten genes discarded. Hsp70 and J-domain sequences were aligned with the HMMER package [80]; for Hsp70 we used the Pfam hidden Markov model profile PF00012, whereas for J-domains we constructed a custom profile. For the HK/RR and ModBC datasets, we picked sequences out of the curated datasets that were not used to train the AttentionDCA models.

In all datasets used to test CLAPP, organisms containing only a single paralog in each family were excluded, since their matching is trivial and would artificially inflate performance metrics. After filtering, there were 213 pairs for Hsp70s and J-domains, 136 pairs for HK/RR and 301 for ModBC.

For the Hsp70 and J-domain families, we also curated a labeled dataset for each family in which sequences were assigned to the subclasses corresponding to the three known *E. coli* chaperone systems [11]. Hsp70 sequences were divided into the subclasses DnaK, HscA, and HscC, whereas J-domain sequences were divided into DnaJ, HscB, and DjlC. To assign these labels, we built HMM profiles for each subclass from high-confidence seed sequences and used *hmmscan* from HM-MER [80] to classify bacterial reference-proteome sequences carrying InterPro identifiers IPR013126 and IPR036869. After this step, sequences were further filtered using their PCA coordinates, with sequences lying within clusters inconsistent with their assigned subclass being removed. These datasets provide an independent biological reference for evaluating whether unsupervised labeling methods recover functionally meaningful subclasses. However, they were not used for paralog matching: the stringent filtering required to retain only high-confidence subclass assignments left too little overlap with the sequence pairs for which genomic proximity could provide a reliable interaction ground truth.

#### S1.2.1 Supplementary tables and figures

**Fig. S1:**
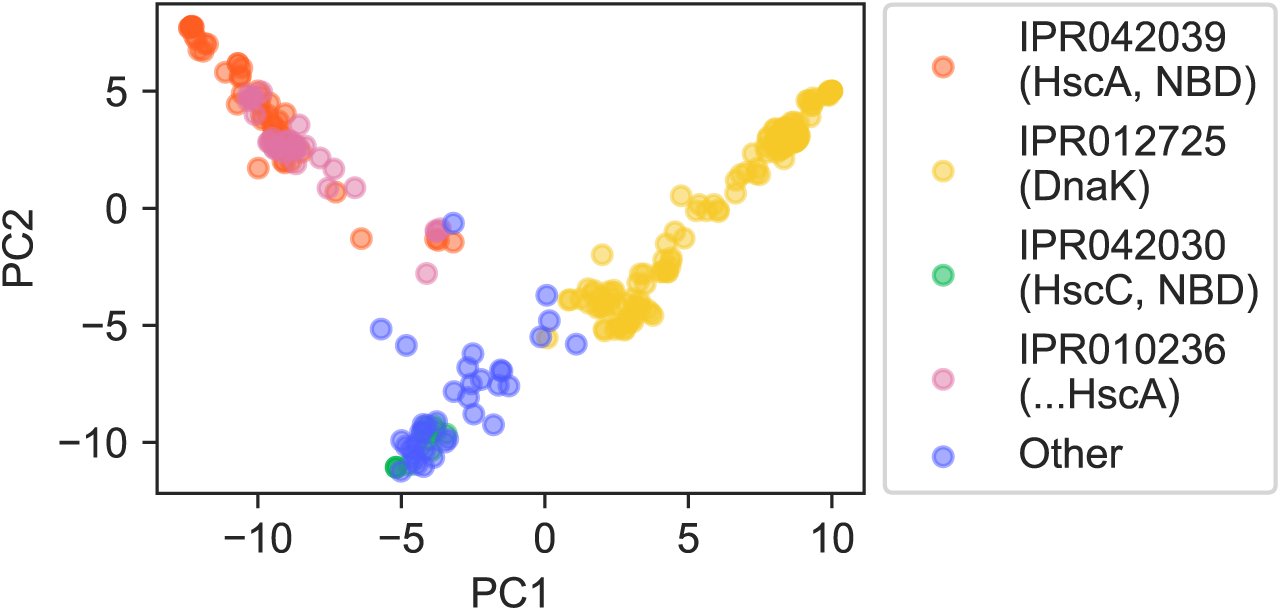
**PCA of Hsp70 test sequences, color-coded according to their InterPro identifiers.**

**Fig. S2:**
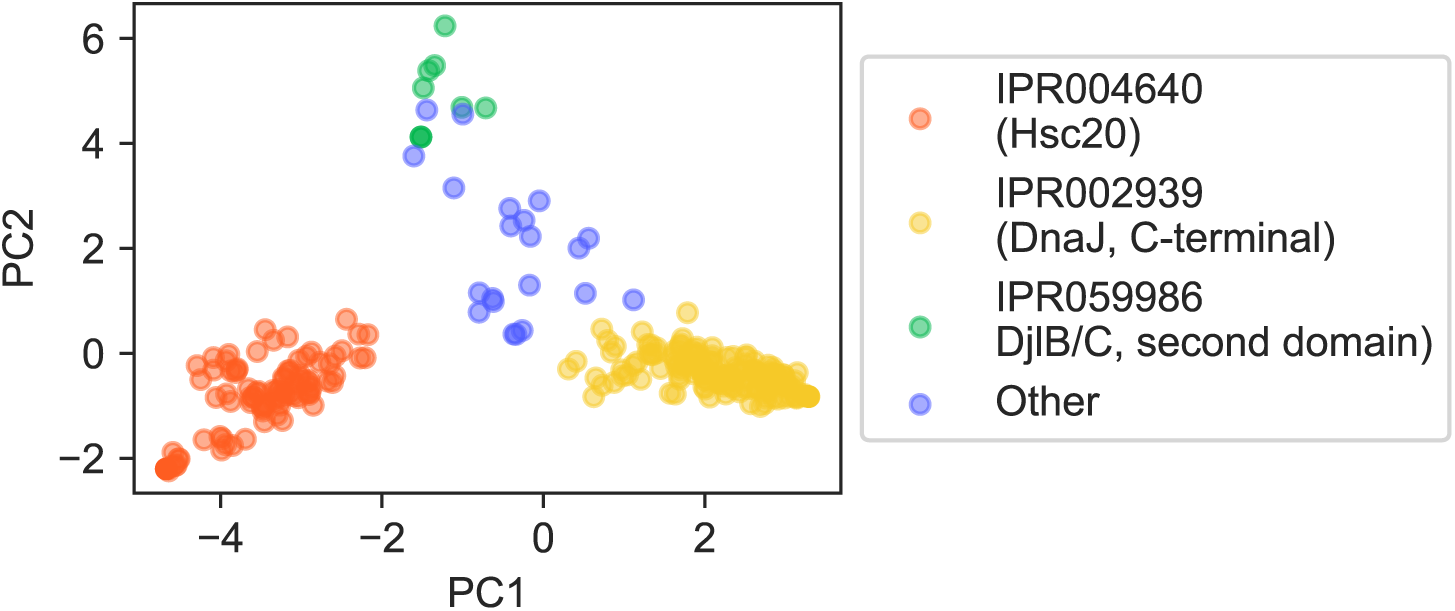
PCA of J-domain sequences to be paired, color-coded according to their InterPro identifiers.

**Fig. S3:**
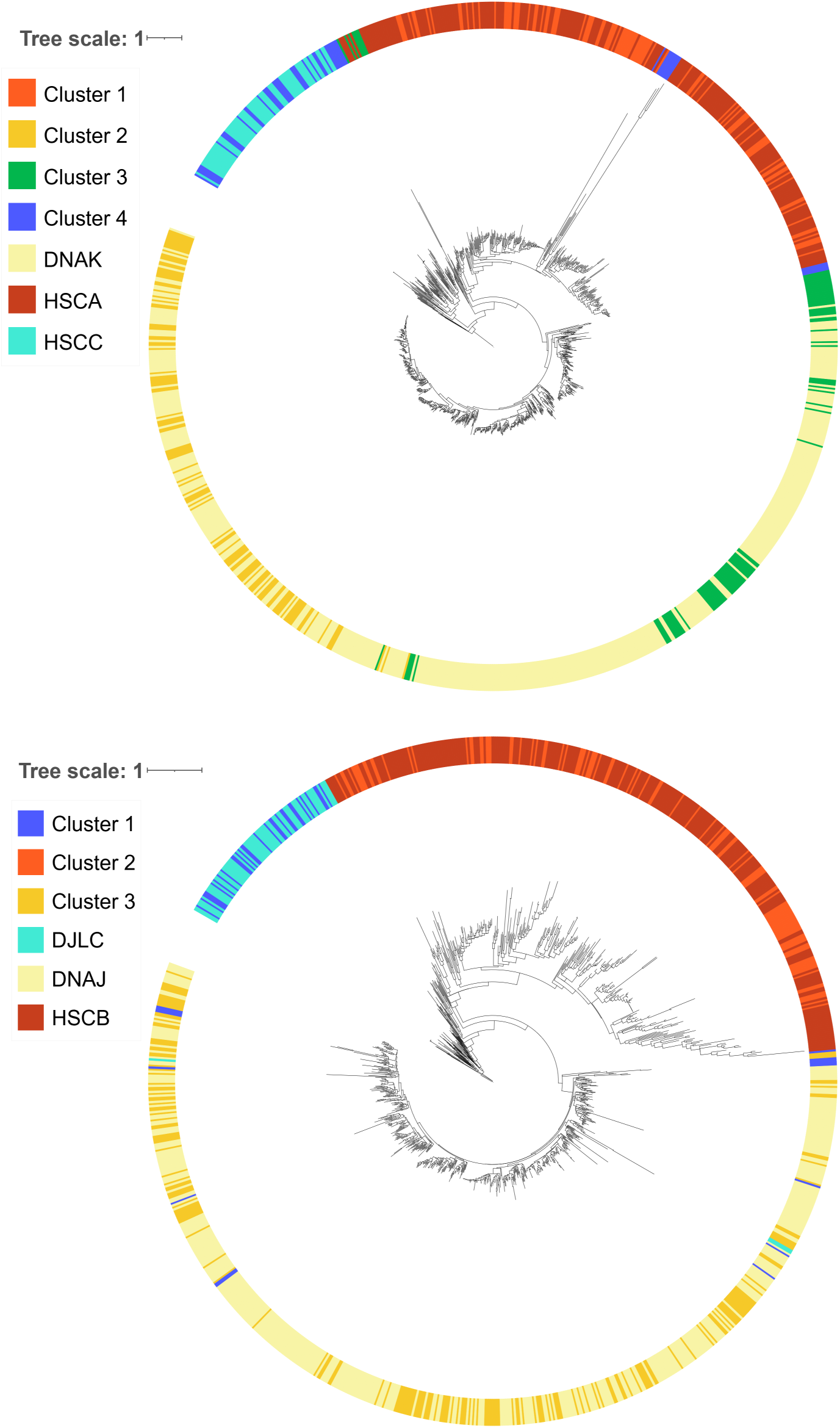
**Phylogenetic trees of Hsp70 and J-domain sequences**, top and bottom panels respectively. Sequences used to construct the trees include Hsp70 (resp. J-domain) sequences labeled using the procedure described in appendix S1.2, as well as sequences labeled using PCA and k-means.

**Fig. S4:**
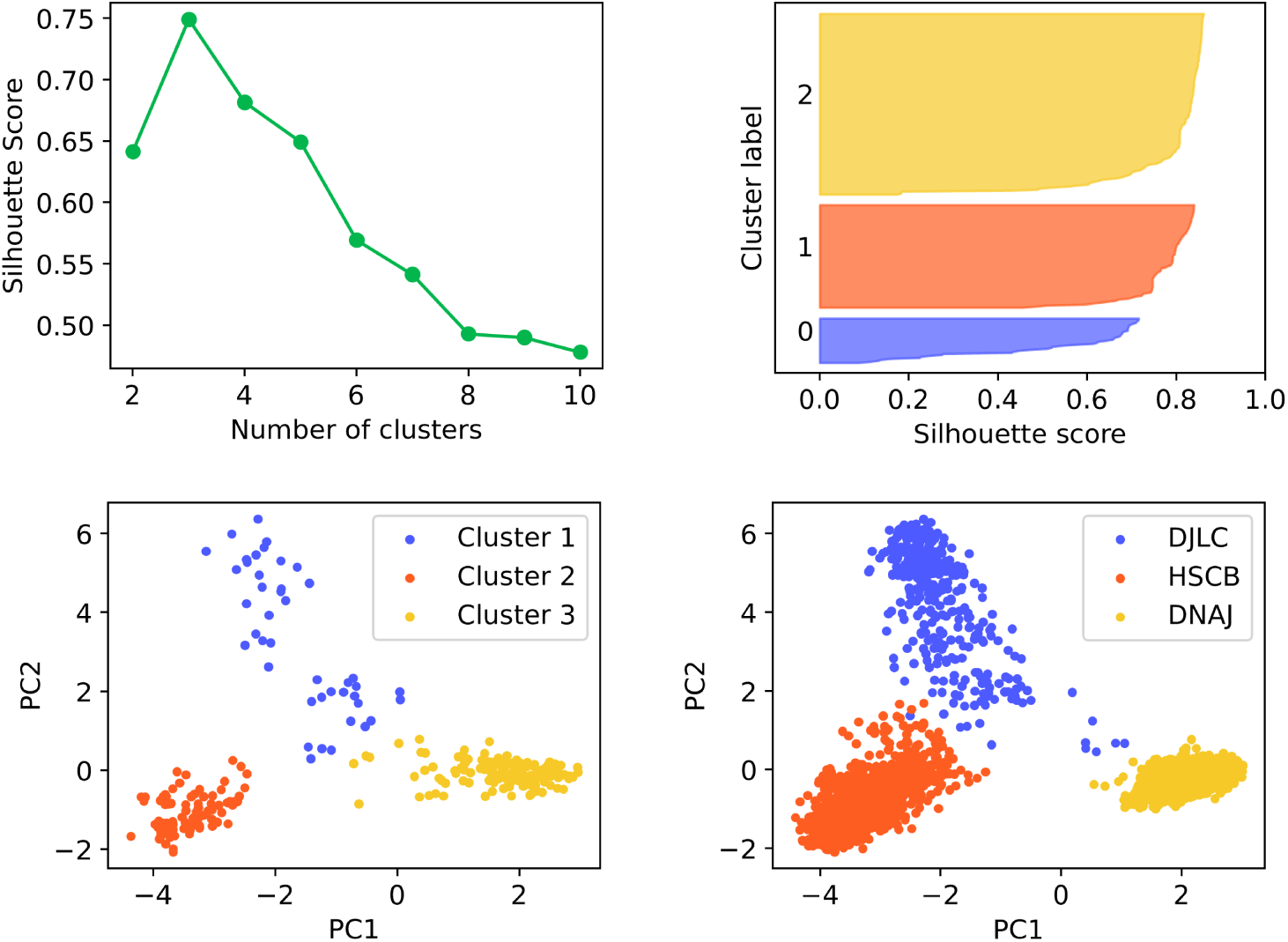
K-means clustering results of J-domain sequences. The top left panel shows the silhouette score as a function of the number of k-means clusters for the first two principal components of the sequences, while the top right panel shows the distribution of silhouette scores for all sequences, divided into three k-means clusters. The bottom left panel shows the PCA of the sequences, color coded according to the k-means clusters. The bottom right panel shows the PCA of a bigger J-domain dataset, color coded according to their known functional subtypes. The k-means clusters correspond to the functional subtypes.

**Fig. S5:**
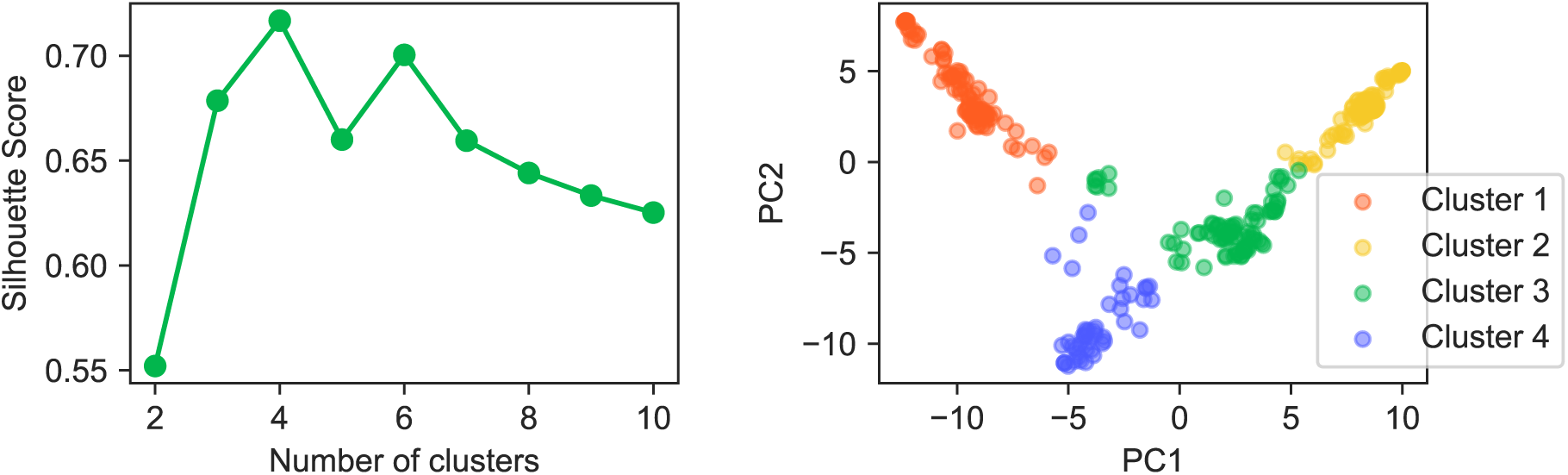
**K-means clustering results of Hsp70 sequences.**The left panel shows the silhouette score as a function of the number of k-means clusters for the first two principal components of the sequences, while the right panel shows the PCA of the sequences, color coded according to the k-means clusters.

**Fig. S6:**
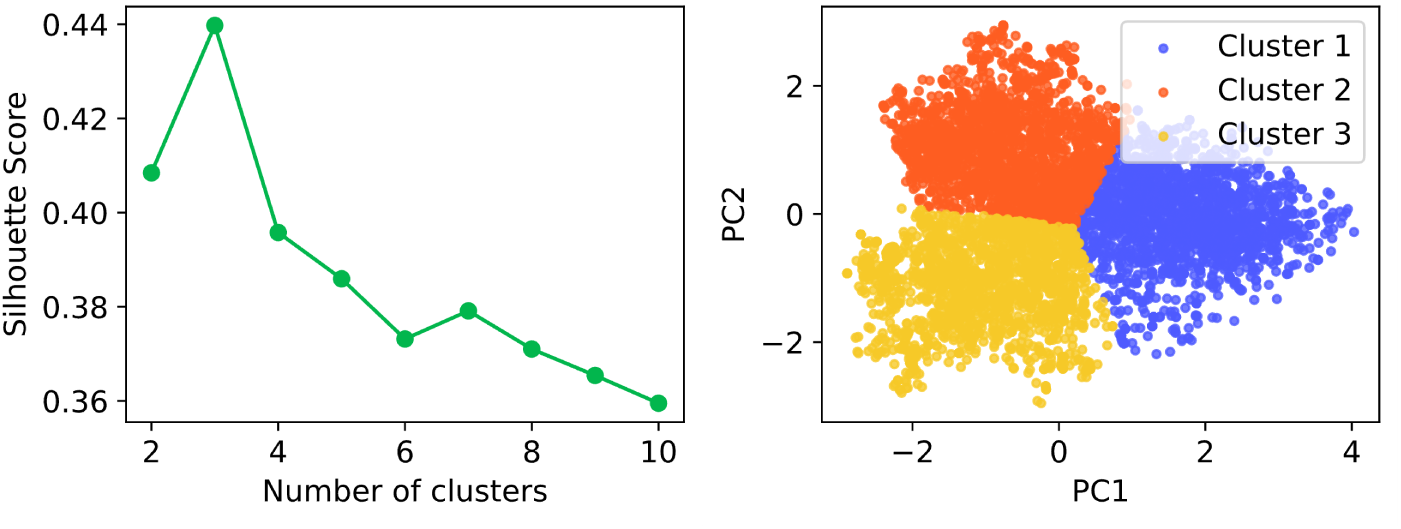
K-means clustering results of HK sequences. The left panel shows the silhouette score as a function of the number of k-means clusters for the first two principal components of the sequences, while the right panel shows the PCA of the sequences, color coded according to the k-means clusters.

**Fig. S7:**
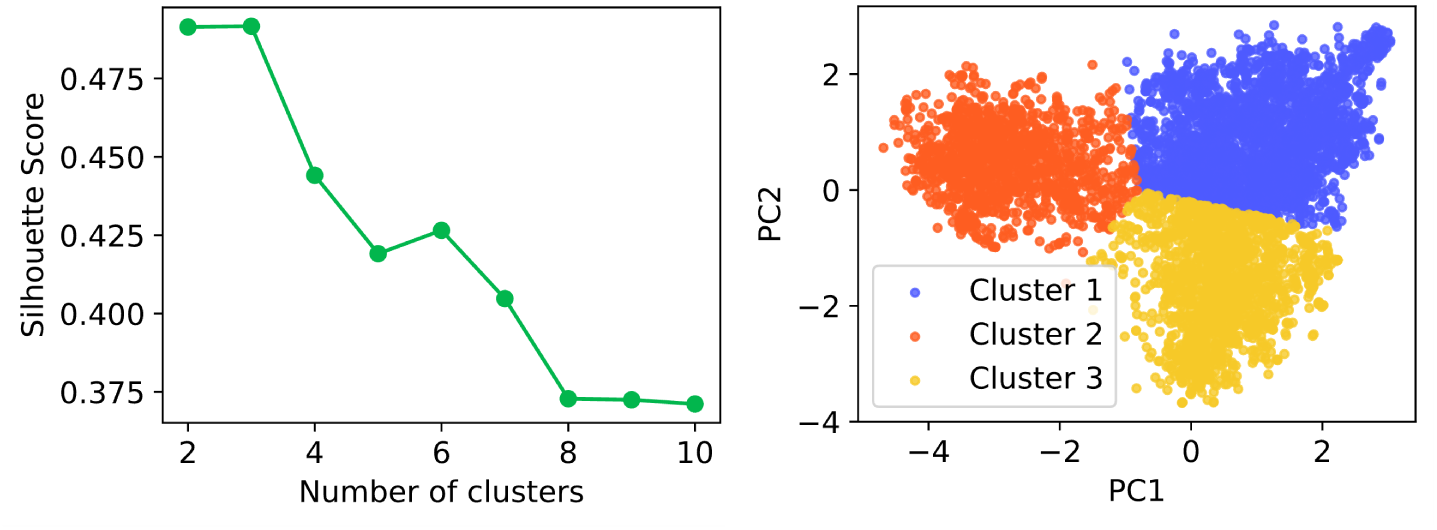
K-means clustering results of RR sequences. The left panel shows the silhouette score as a function of the number of k-means clusters for the first two principal components of the sequences, while the right panel shows the PCA of the sequences, color coded according to the k-means clusters.

**Fig. S8:**
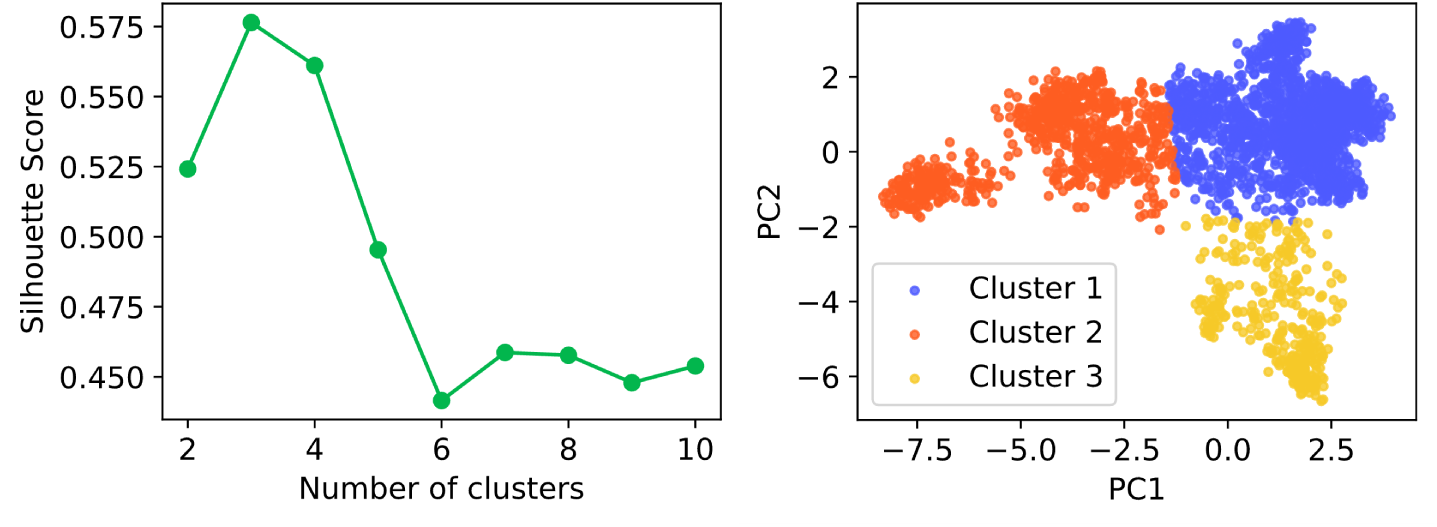
K-means clustering results of ModC sequences. The left panel shows the silhouette score as a function of the number of k-means clusters for the first two principal components of the sequences, while the right panel shows the PCA of the sequences, color coded according to the k-means clusters.

**Fig. S9:**
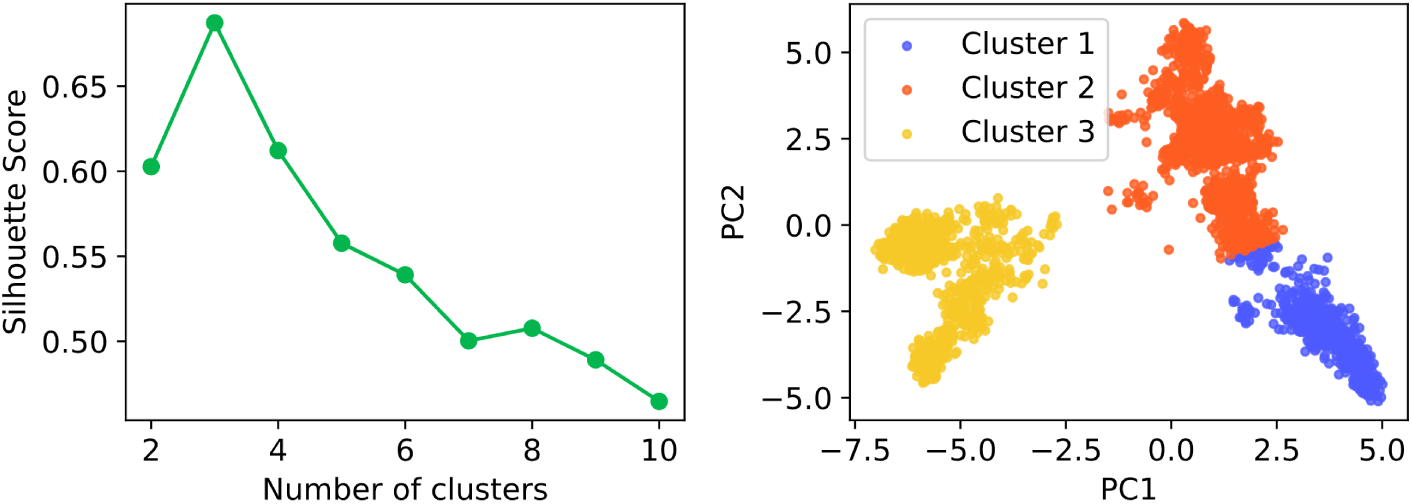
K-means clustering results of ModB sequences. The left panel shows the silhouette score as a function of the number of k-means clusters for the first two principal components of the sequences, while the right panel shows the PCA of the sequences, color coded according to the k-means clusters.

**Fig. S10:**
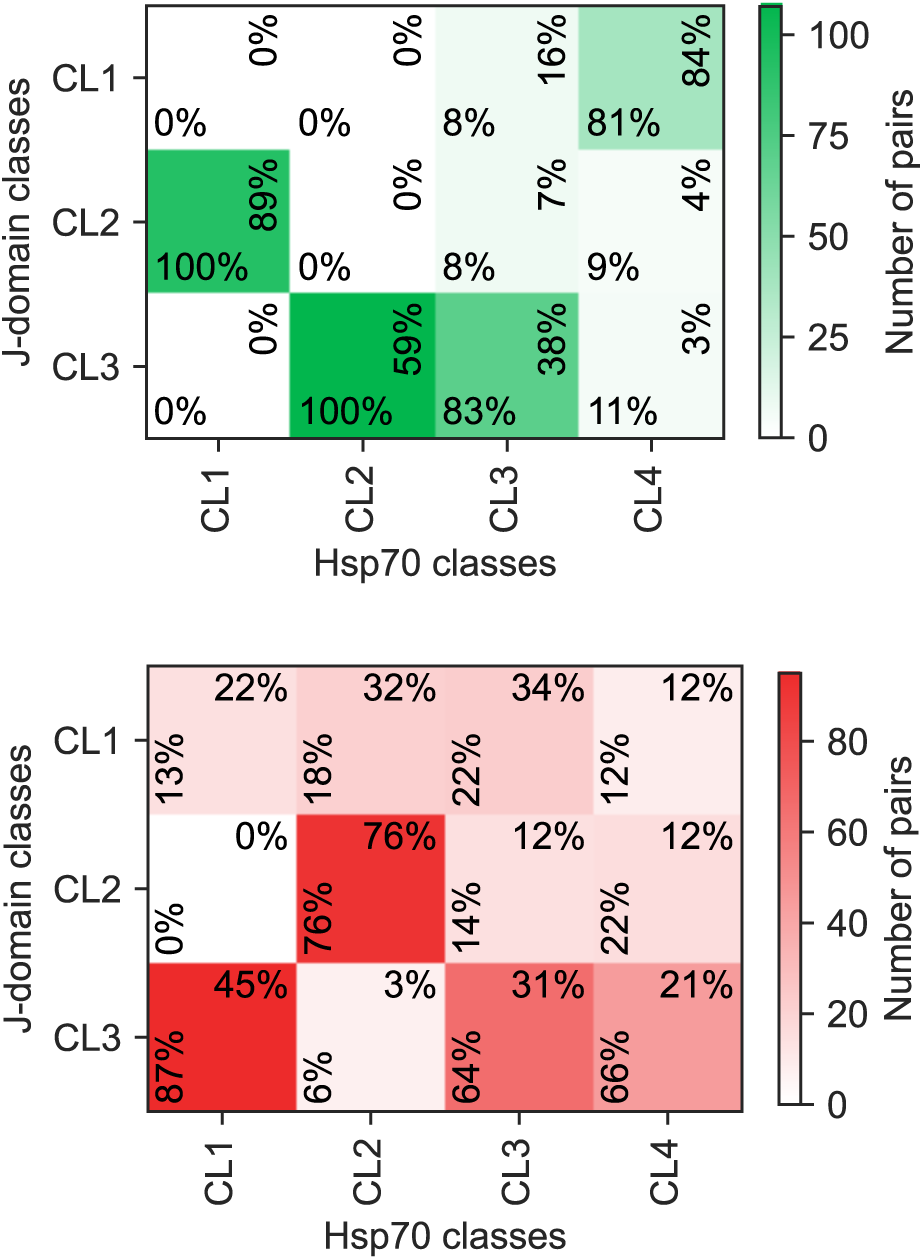
Distribution of J-domain and Hsp70 within-species pairs in the possible cluster combinations. The upper panel shows the distribution of correct pairs according to genomic proximity, while the lower panel shows the distribution of all within-species wrong pairs. Percentages written horizontally (resp. vertically) represent the fraction of Hsp70s (resp. J-domains) in each category. In the final dataset used for testing, we only keep J-domain and Hsp70 pairs of the type CL1-CL4, CL2-CL1 and CL3-CL2.

**Fig. S11:**
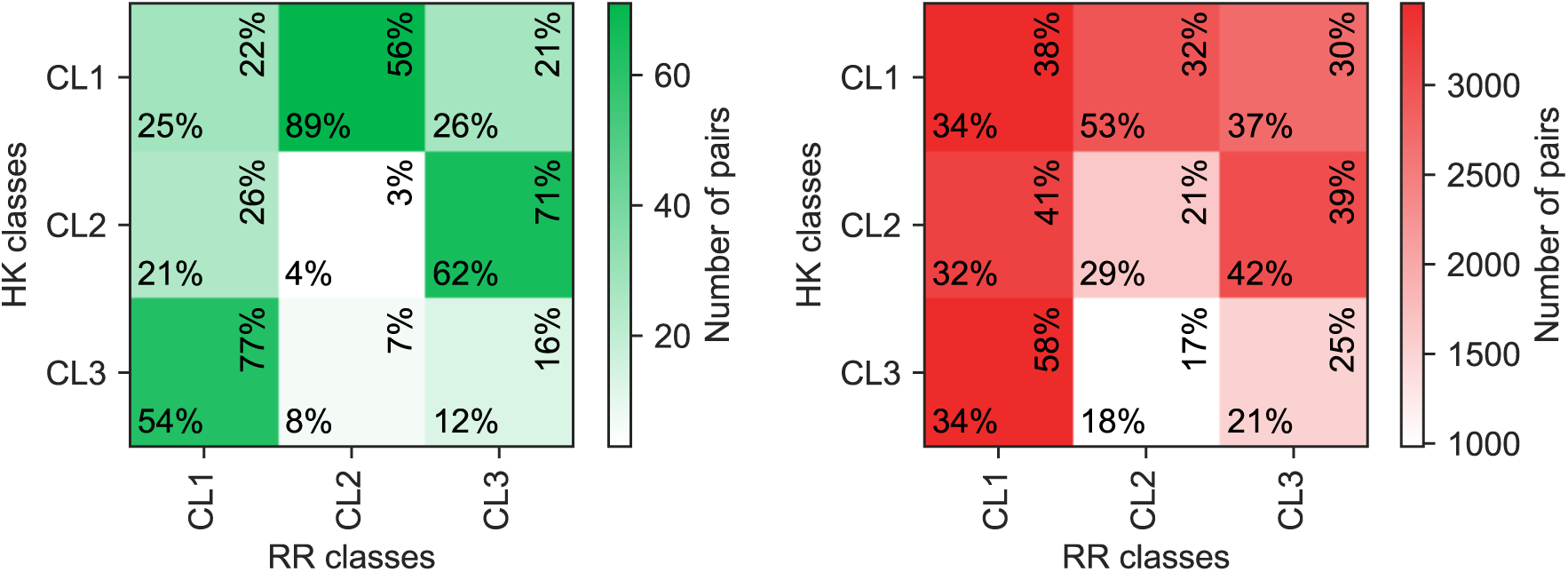
Distribution of HK and RR within-species pairs in the possible cluster combinations. The upper panel shows the distribution of correct pairs according to genomic proximity, while the lower panel shows the distribution of all within-species wrong pairs. Percentages written horizontally (resp. vertically) represent the fraction of RR (resp. HK) in each category. In the final dataset used for testing, we only keep pairs of the type CL1-CL2, CL2-CL3 and CL3-CL1.

**Fig. S12:**
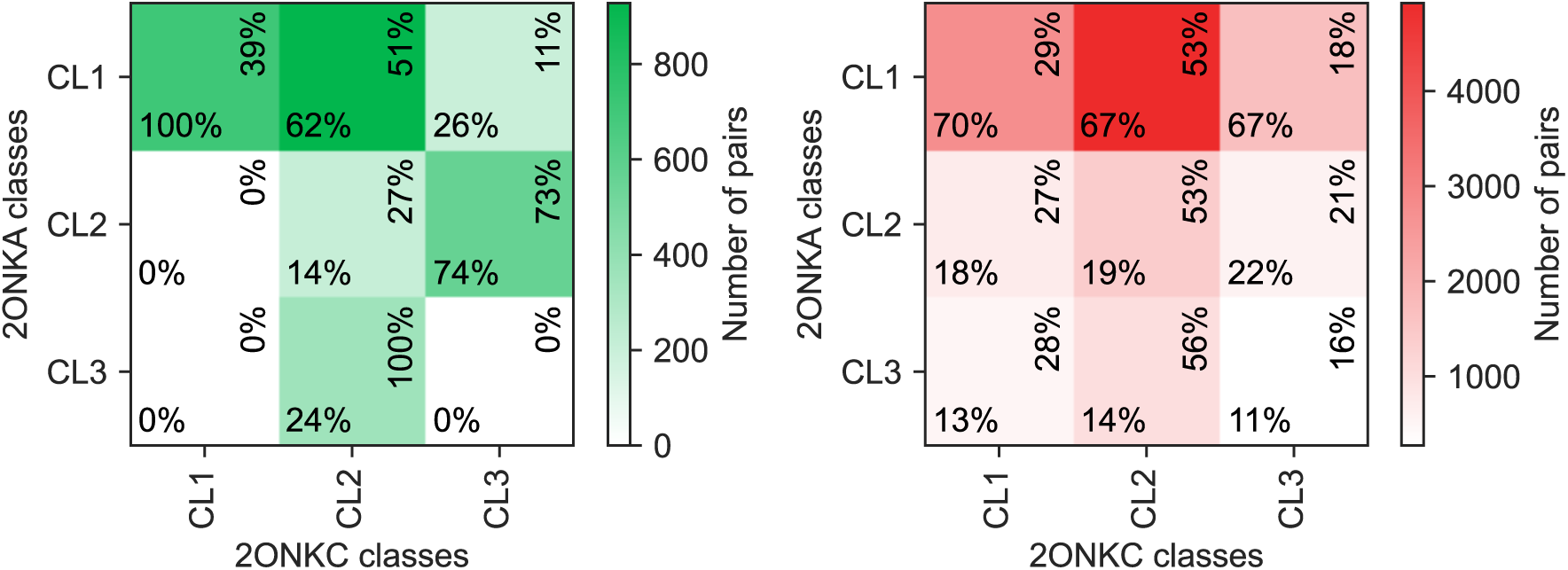
Distribution of ModC and ModB within-species pairs in the possible cluster combinations. The upper panel shows the distribution of correct pairs according to genomic proximity, while the lower panel shows the distribution of all within-species wrong pairs. Percentages written horizontally (resp. vertically) represent the fraction of ModB (resp. ModC) in each category. In the final dataset used for testing, we only keep pairs of the type CL1-CL1, CL2-CL3 and CL3-CL2.

**Fig. S13:**
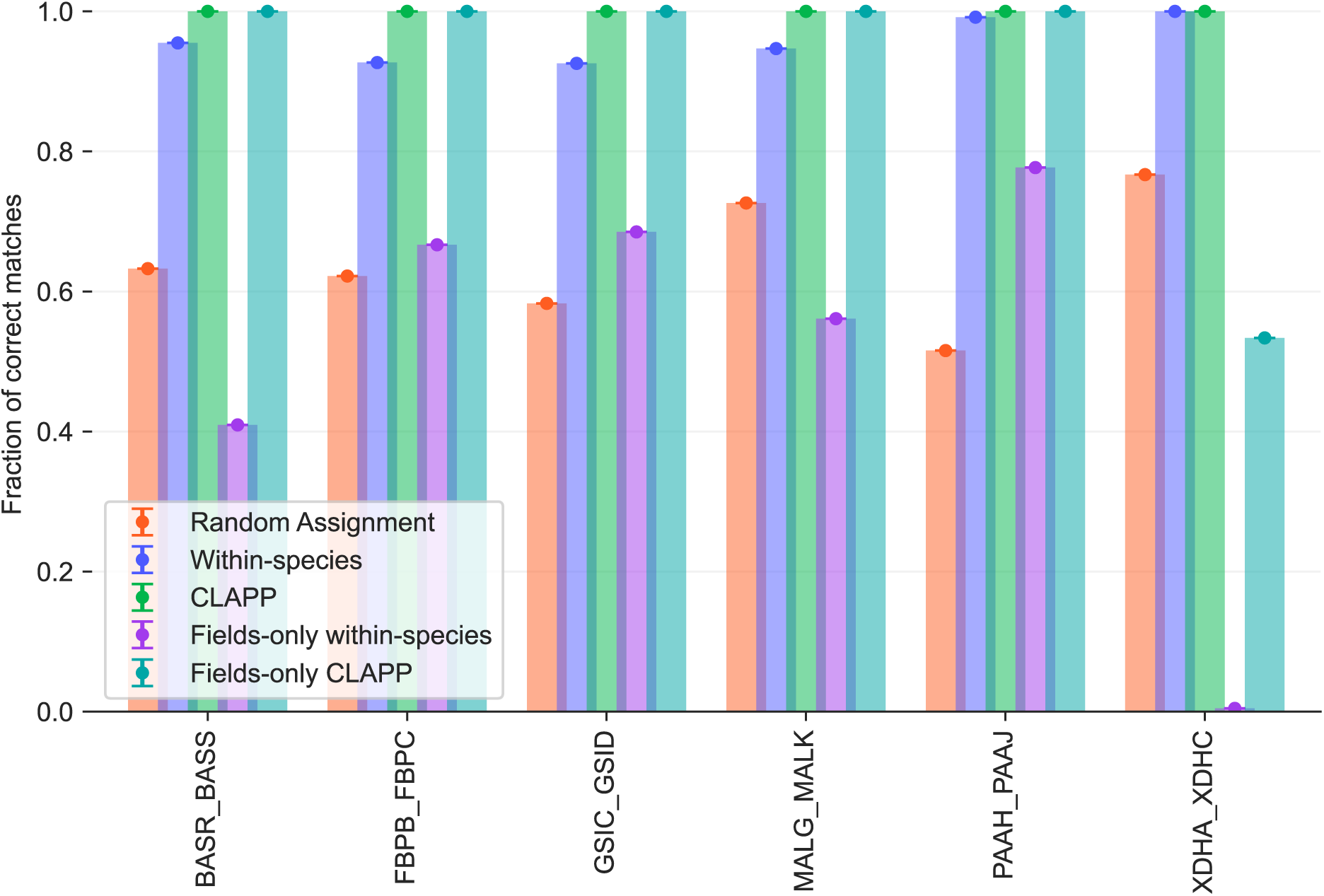
**Paralog matching performance of CLAPP using different sequence-based interaction scores**, on concatenated MSAs from [36] and filtered to enforce one-to-one interactions between classes. Fraction of correct matches obtained using AttentionDCA and fields-only scores after class-level pooling. Performance is compared to the random assignment of within-species pairs. Sequences are divided into classes using k-means, and class labels are used to construct the pooling matrix. Results are averaged over five random initializations, with error bars indicating one standard deviation.

**Fig. S14:**
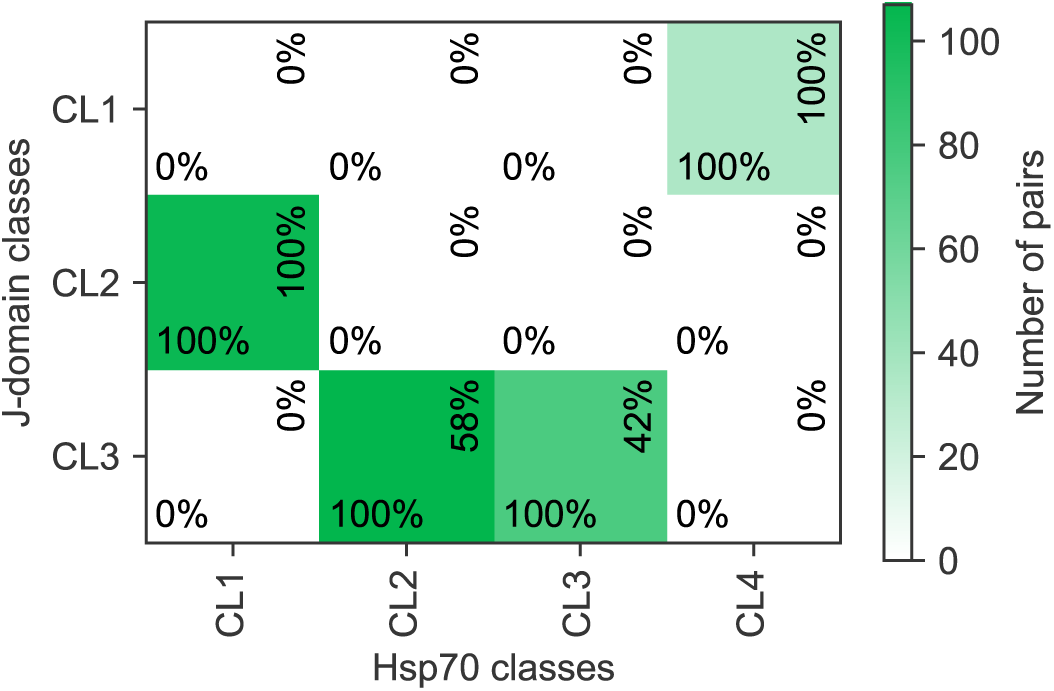
Distribution of J-domain and Hsp70 within-species pairs in the possible cluster combinations, after phylogeny-informed relabeling, according to genomic proximity. Percentages written horizontally (resp. vertically) represent the fraction of Hsp70s (resp. J-domains) in each category. In the final dataset used for testing, we merge Hsp70 classes CL2 and CL3.

**Fig. S15:**
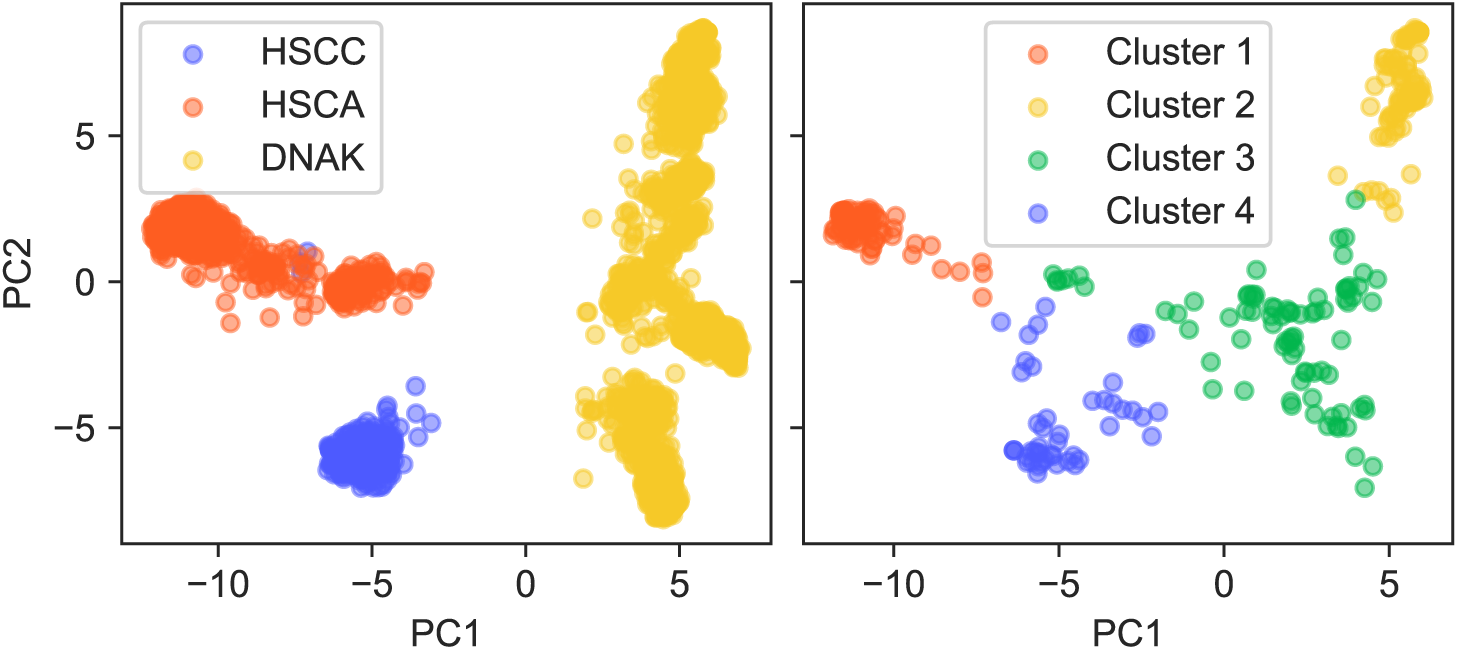
K-means clustering results of Hsp70 sequences. The left panel shows the PCA of Hsp70 sequences color-coded using architectural labels, while the right panel shows Hsp70 sequences used for testing, projected onto the principal components of labeled sequences and color coded according to the k-means clusters.

**Fig. S16:**
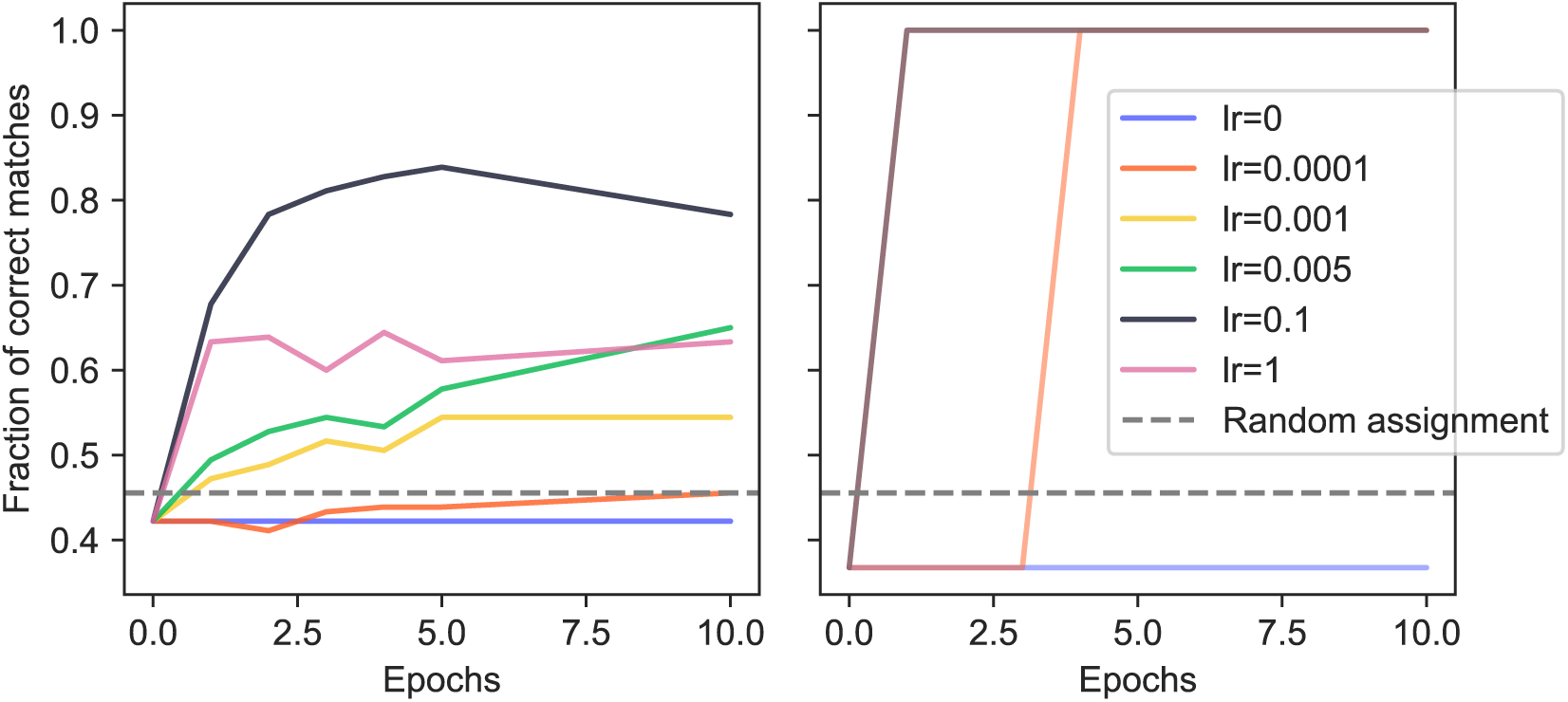
**Fraction of correct matches using the AttentionDCA score within-species (left) and with pooling matrix (CLAPP, right) on the HK/RR dataset, as a function of the number of training epochs, for different learning rate values**. Sequences are divided into clusters using k-means and cluster labels are used for the pooling matrix. Performance is compared to the random assignment of within-species pairs. Median over 5 random initializations. Default learning rate value used when not specified is 0.005. As expected, when the learning rate is zero, performance remains constant at the random-assignment baseline. Decreasing the learning rate delays convergence, requiring more epochs to achieve the same performance, whereas excessively large learning rates prevent the model from reaching optimal performance.

**Fig. S17:**
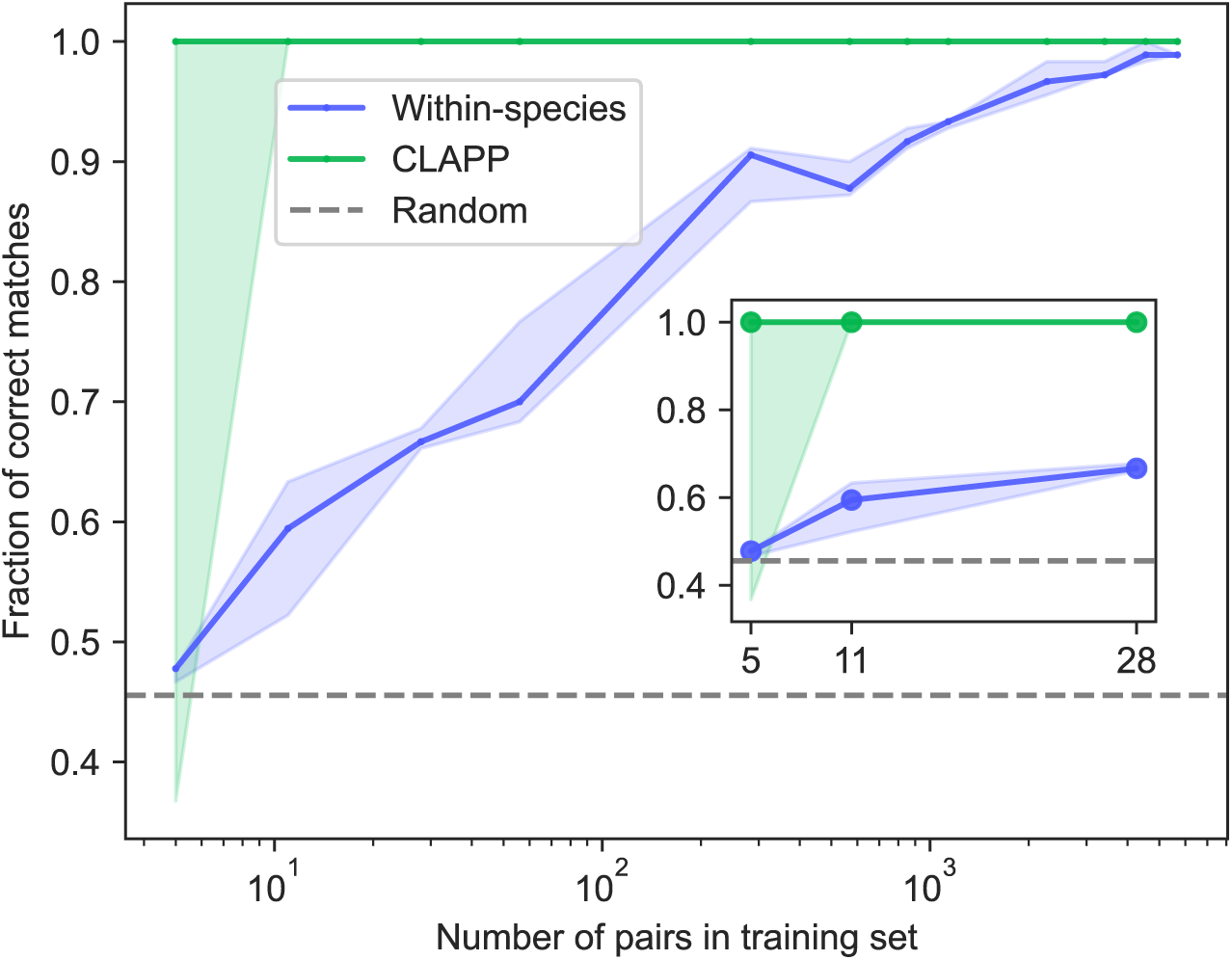
**Fraction of correct matches using the AttentionDCA score within-species and with pooling matrix (CLAPP), as a function of the number of pairs in the training set for the HK/RR dataset**. The AttentionDCA model is trained on different numbers of only correct pairs (on the x-axis), which is why performance is higher than in the left panel of Fig. 7. Inset shows performance for 5-28 pairs in the alignment used to train AttentionDCA, while main panels show performance starting from 5 pairs in log scale. Sequences are divided into clusters using k-means and cluster labels are used for the pooling matrix. Performance is compared to the random assignment of within-species pairs. Matching performed for 5 random initializations, solid line is the median and shaded area corresponds to interquartile range. For each random initialization, a new training MSA is selected with only correct pairs, such that all pairs in an organism are either included or excluded. The actual number of training pairs therefore slightly oscillates between random seeds.

**Fig. S18:**
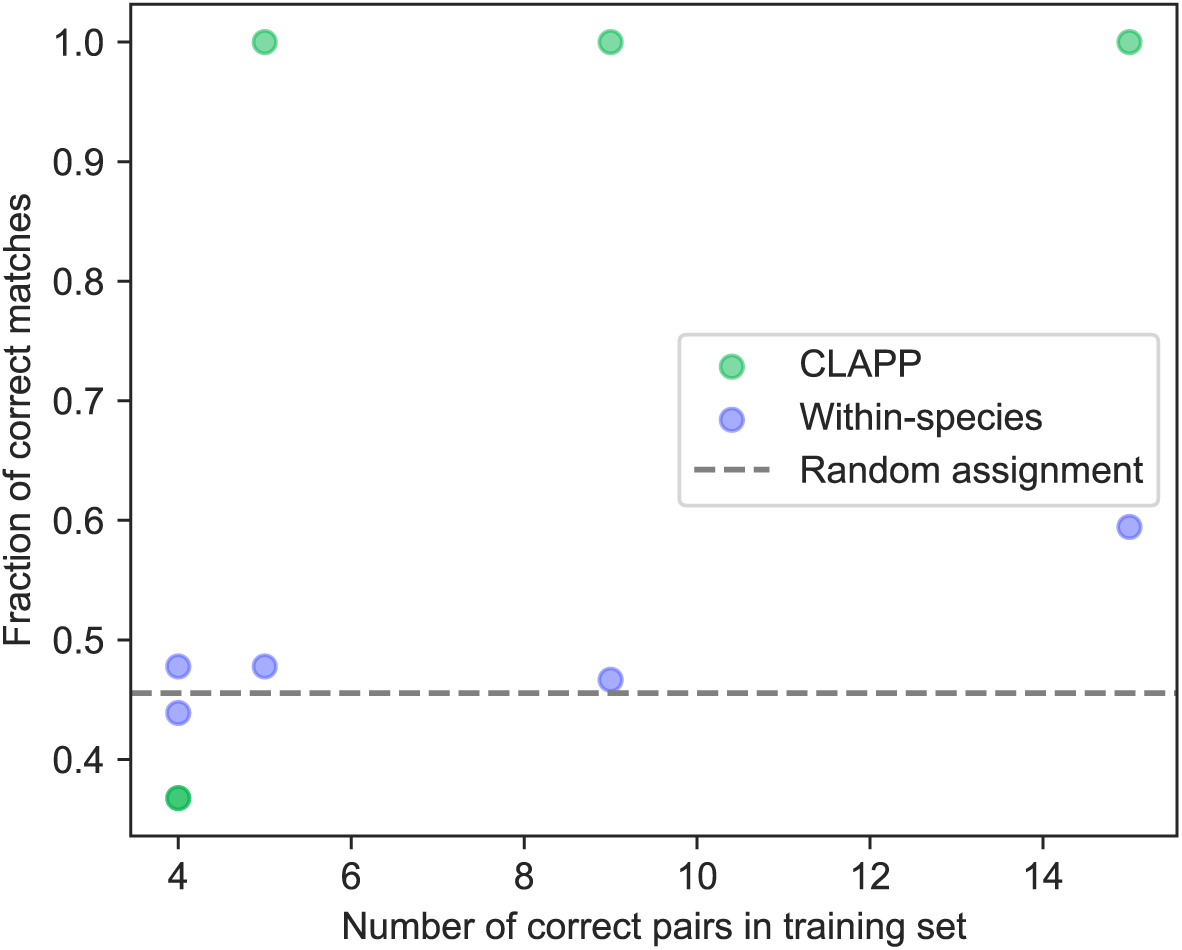
Fraction of correct matches as a function of the number of pairs in the training set for the HK/RR dataset and low number of correct pairs in the training set. Each random initialization result is shown here as a point.

**Fig. S19:**
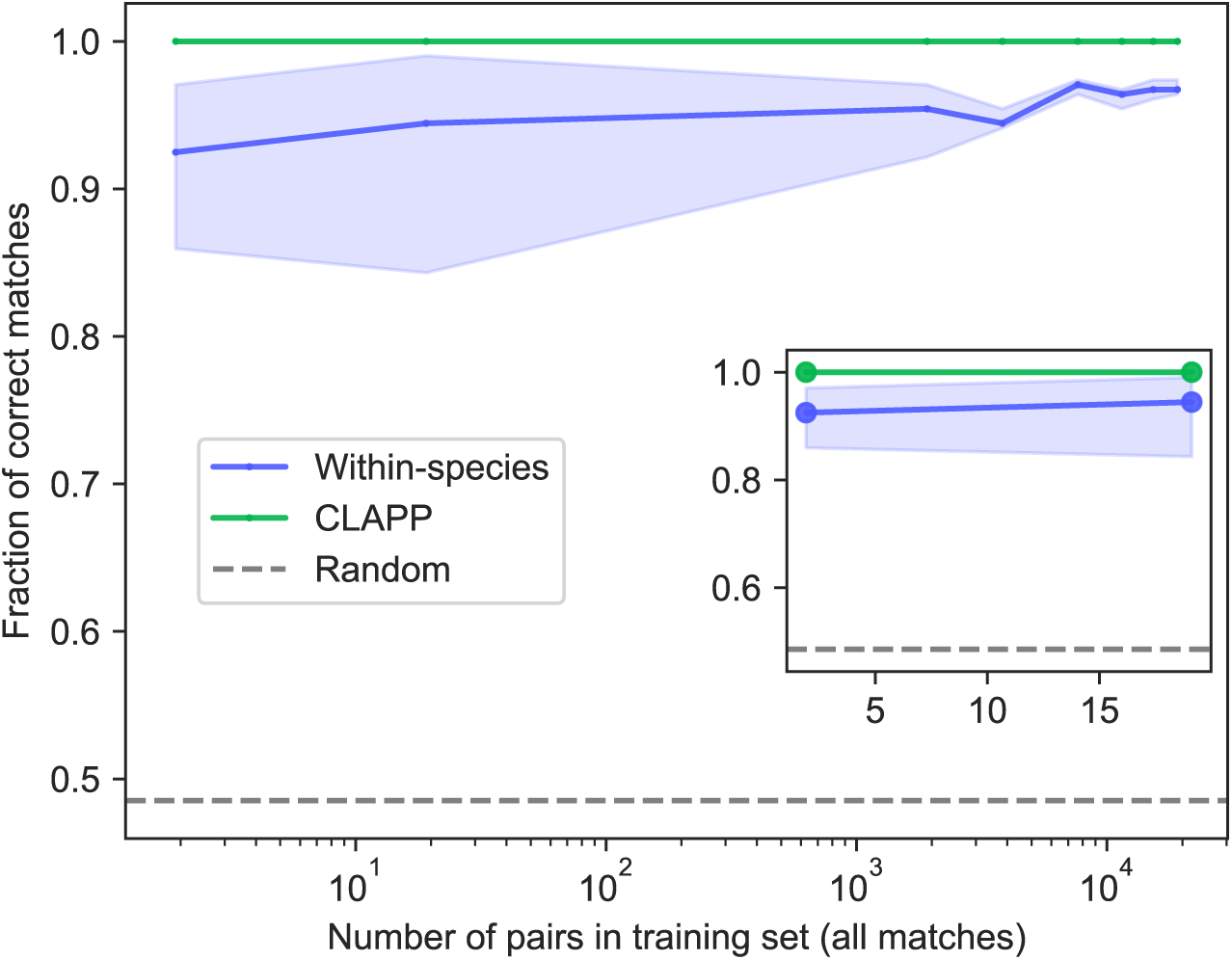
**Fraction of correct matches using the AttentionDCA score within-species and with pooling matrix (CLAPP), as a function of the number of pairs in the training set for the J-domain/Hsp70 dataset**. The AttentionDCA model is trained on all matches assembled from different numbers of all correct pairs, on the x-axis. Inset shows performance for 2-19 pairs in the alignment used to train AttentionDCA, while main panels show performance starting from 2 pairs in log scale. Sequences are divided into clusters using k-means and cluster labels are used for the pooling matrix. Performance is compared to the random assignment of within-species pairs. Matching performed for 5 random initializations, solid line is the median and shaded area corresponds to interquartile range.

**Fig. S20:**
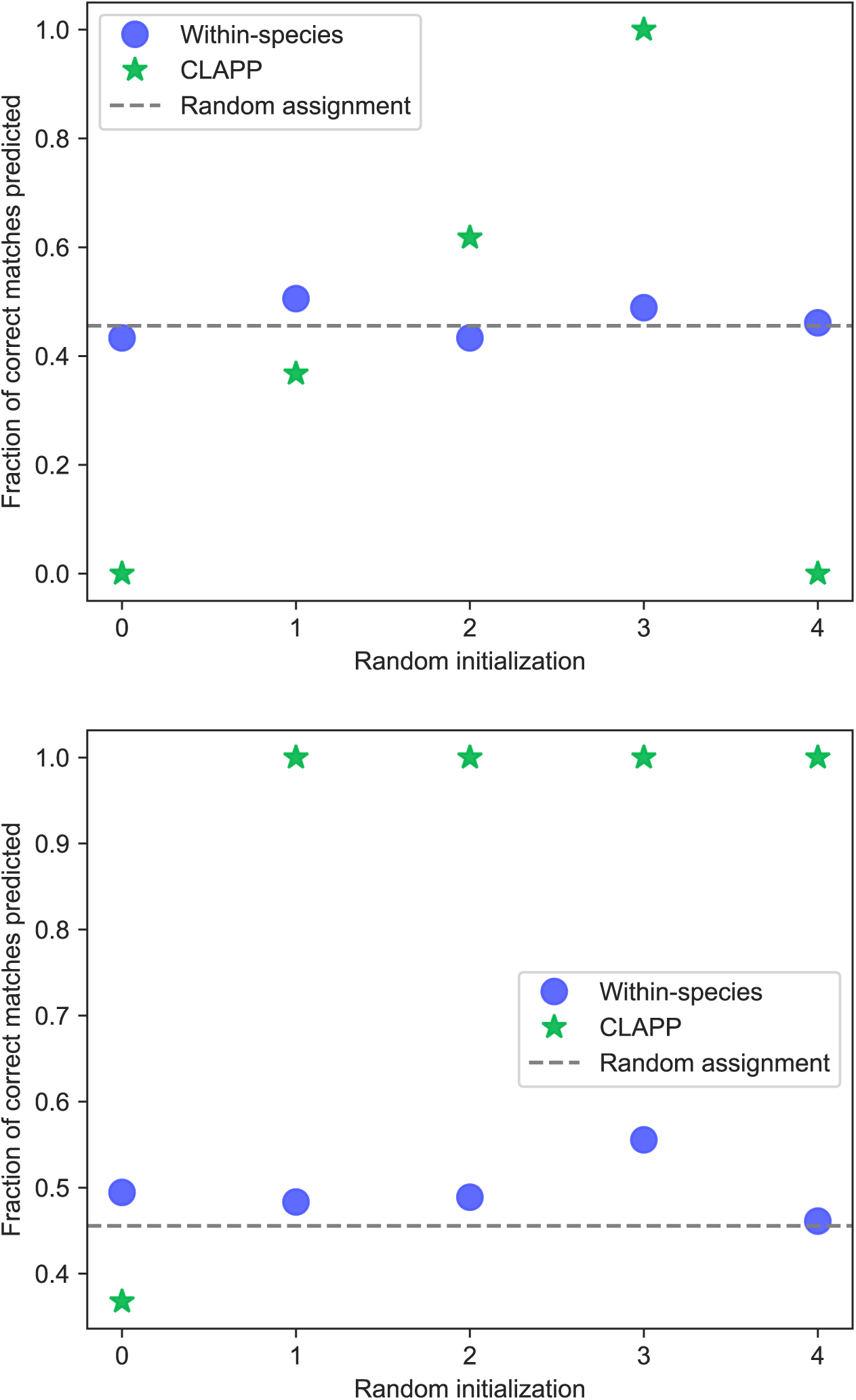
Fraction of correct matches using the AttentionDCA score within-species and with pooling matrix (CLAPP), for different initializations. The AttentionDCA model is trained for 0 epochs (top) and 1 epoch (bottom) on HK/RR datasets with no correct pairs

**Fig. S21:**
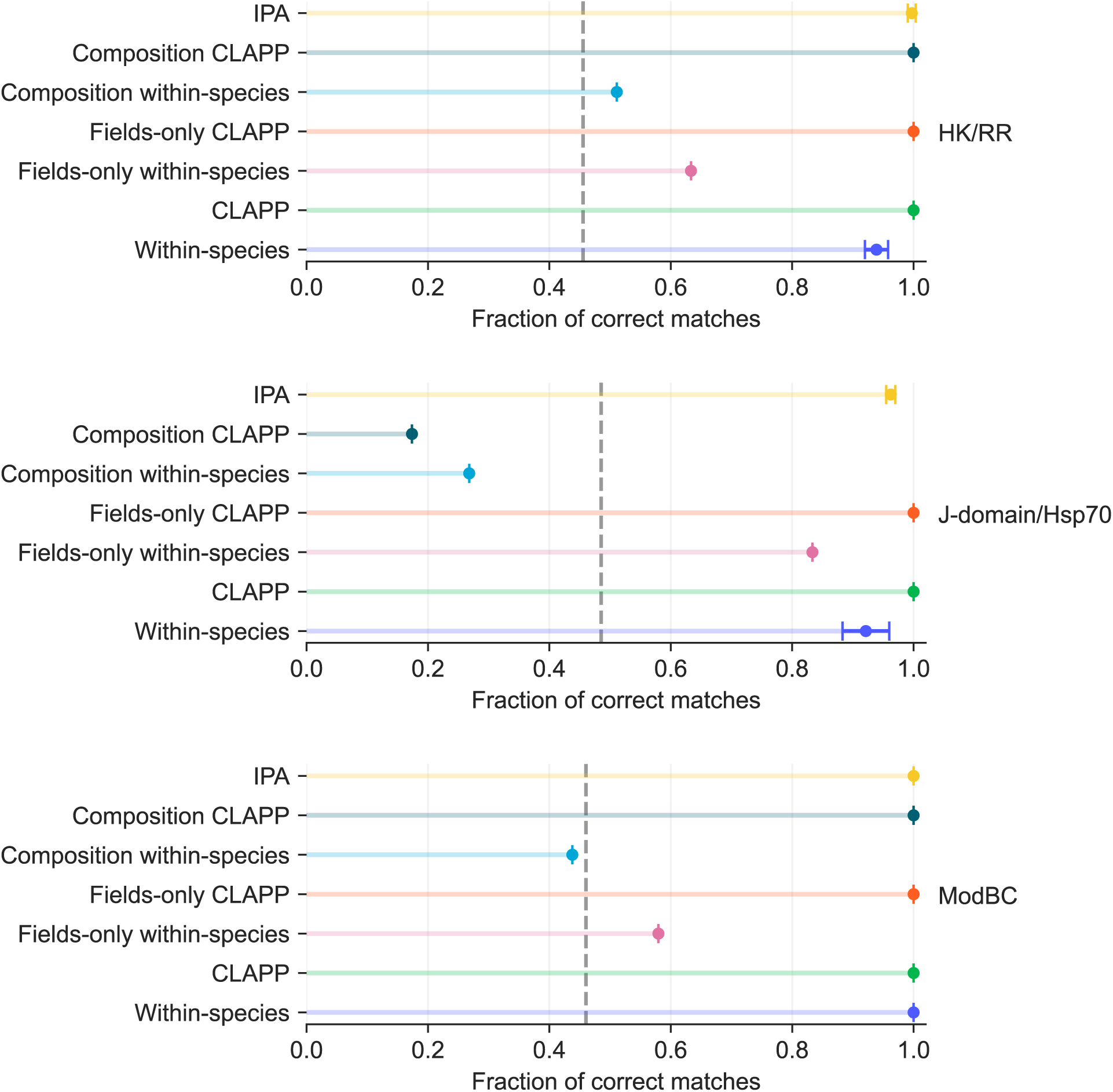
Paralog matching performance of AttentionDCA, fields-only and composition scores within-species and with pooling matrix (CLAPP). Sequences are divided into clusters using k-means and cluster labels are used for the pooling matrix. Results are averaged over five random realizations, with error bars indicating one standard deviation.

**Fig. S22:**
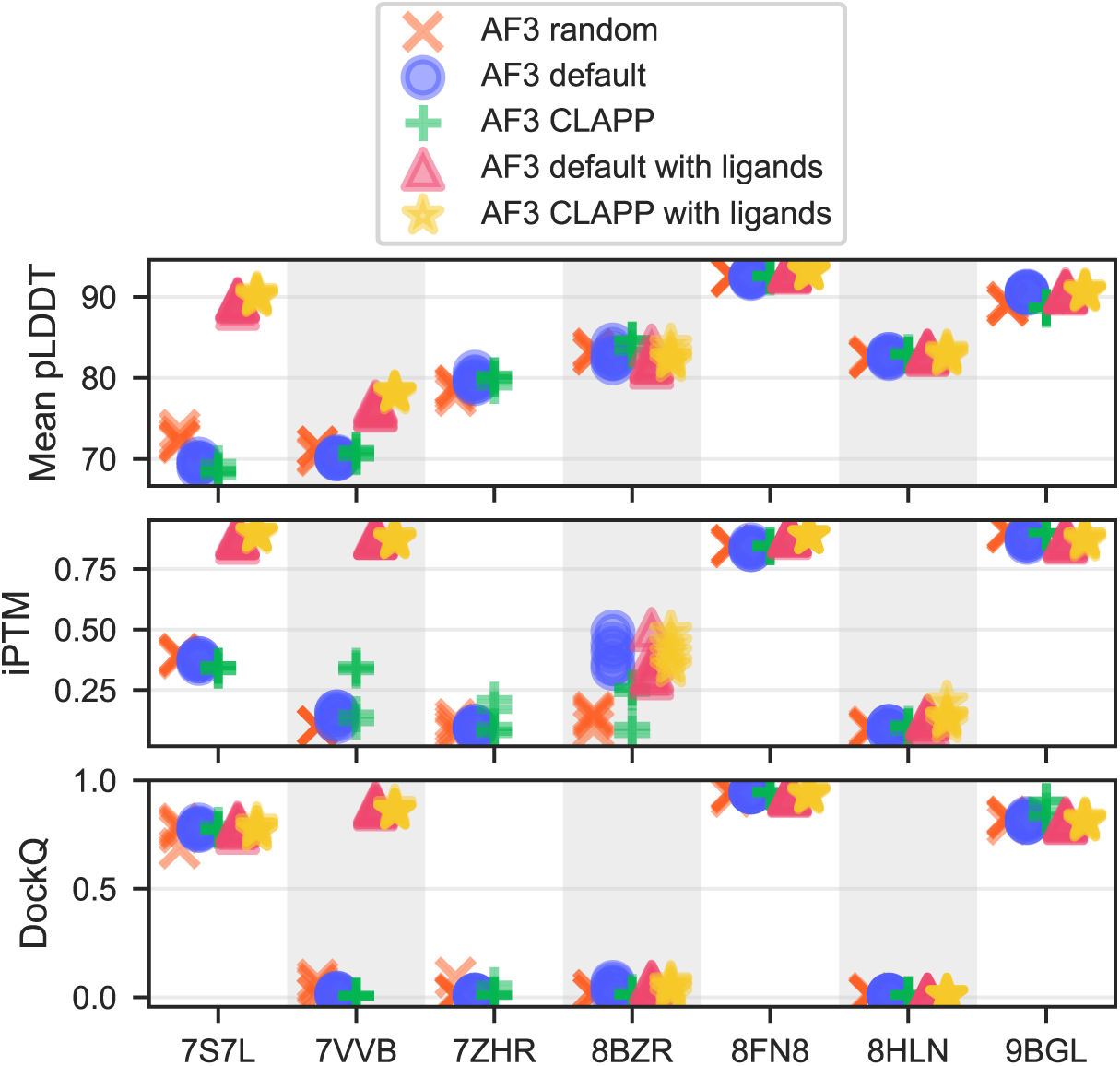
Comparison of MSA pairing methods for complex structure prediction with AlphaFold 3. For each experimental structure, five models were generated for each method. The top, middle, and bottom panels show mean pLDDT, ipTM, and DockQ scores, respectively. Higher values indicate better predictions for all three metrics.

**Table S1:**
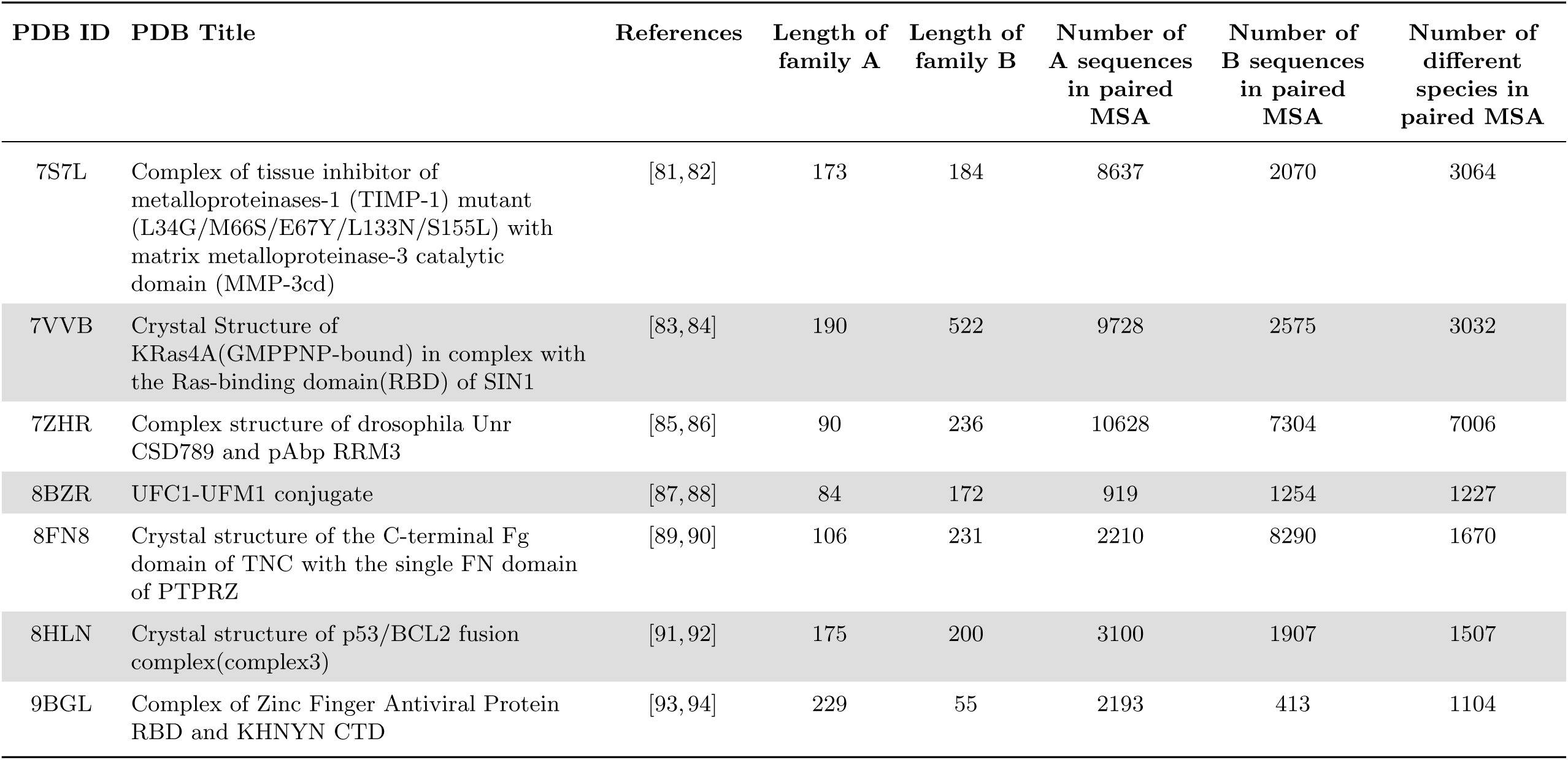
PDB and paired MSA information for eukaryotic complex structure prediction.

| PDB ID | PDB Title | References | Length of family A | Length of family B | Number of A sequences in paired MSA | Number of B sequences in paired MSA | Number of different species in paired MSA |
| --- | --- | --- | --- | --- | --- | --- | --- |
| 7S7L | Complex of tissue inhibitor of metalloproteinases-1 (TIMP-1) mutant (L34G/M66S/E67Y/L133N/S155L) with matrix metalloproteinase-3 catalytic domain (MMP-3cd) | [81, 82] | 173 | 184 | 8637 | 2070 | 3064 |
| 7VVB | Crystal Structure of KRas4A(GMPPNP-bound) in complex with the Ras-binding domain(RBD) of SIN1 | [83, 84] | 190 | 522 | 9728 | 2575 | 3032 |
| 7ZHR | Complex structure of drosophila Unr CSD789 and pAbp RRM3 | [85, 86] | 90 | 236 | 10628 | 7304 | 7006 |
| 8BZR | UFC1-UFM1 conjugate | [87, 88] | 84 | 172 | 919 | 1254 | 1227 |
| 8FN8 | Crystal structure of the C-terminal Fg domain of TNC with the single FN domain of PTPRZ | [89, 90] | 106 | 231 | 2210 | 8290 | 1670 |
| 8HLN | Crystal structure of p53/BCL2 fusion complex(complex3) | [91, 92] | 175 | 200 | 3100 | 1907 | 1507 |
| 9BGL | Complex of Zinc Finger Antiviral Protein RBD and KHNYN CTD | [93, 94] | 229 | 55 | 2193 | 413 | 1104 |

## Notes

### Competing Interest Statement

The authors have declared no competing interest.

### Summary of Updates

The schematic representation of the workflow in Figure 1 has been revised for clearness.Sections 3.2-3.5 were added, with corresponding figures. Section 3.4 has been partly moved to the supplementary materials.

### Funding

- Swiss National Science Foundation: IC00I0-227688
- European REA, Marie Skłodowska-Curie Actions: 101131463

